# Inhibition of tumor growth by a novel engineered chimeric toxin that cleaves activated mutant and wild-type RAS

**DOI:** 10.1101/2019.12.17.880187

**Authors:** Vania Vidimar, Greg L. Beilhartz, Minyoung Park, Marco Biancucci, Matthew B. Kieffer, David R. Gius, Roman A. Melnyk, Karla J. F. Satchell

**Affiliations:** Department of Microbiology and Immunology, Northwestern University, Feinberg School of Medicine, Chicago, IL 60611, USA; Department of Radiation Oncology, Northwestern University, Feinberg School of Medicine, Chicago, IL 60611, USA; Robert H. Lurie Comprehensive Cancer Research Center, Northwestern University, Feinberg School of Medicine, Chicago, IL 60611, USA; Program in Molecular Medicine, The Hospital for Sick Children, Toronto, ON M5G 0A4, Canada; Department of Biochemistry, University of Toronto, Toronto, Canada

**Author notes:** GSK Vaccines, Rockville, MD 20850, USA. St. Jude Children’s Research Hospital, Memphis, TN 38105, USA. Co-senior authors (Roman A. Melnyk, Karla J. F. Satchell). Lead contact: Karla Satchell.

## Abstract

Despite nearly four decades of effort, broad inhibition of oncogenic RAS using small molecule approaches has proven to be a major challenge. Here we describe the development of a novel pan-RAS biologic inhibitor comprised of the RAS-RAP1-specific endopeptidase fused to the protein delivery machinery of diphtheria toxin. We show that this engineered chimeric toxin irreversibly cleaves and inactivates intracellular RAS at low picomolar concentrations terminating downstream signaling in receptor-bearing cells. Further, we demonstrate *in vivo* target engagement and reduction of tumor burden in three mouse xenograft models driven by either wild-type or mutant *RAS*. Intracellular delivery of a potent anti-RAS biologic through a receptor-mediated mechanism represents a promising new approach to developing RAS therapeutics against a broad array of cancers.

**Significance:** RAS oncoproteins have long been considered among the most elusive drug targets in cancer research. At issue is the lack of accessible drug binding sites and the extreme affinity for its GTP substrate. Covalent inhibitors against the KRAS G12C mutant have shown early clinical promise, however, targeting the other oncogenic RAS mutants across three RAS isoforms has proven challenging. Inhibition of activated wild-type RAS in the absence of canonical *RAS* mutations is also highly desirable in certain tumors. Here, we demonstrate delivery of an extremely potent pan-RAS and RAP1 cleaving enzyme in therapeutic quantities to specific receptor-bearing cells *in vitro* and *in vivo*. We aim to advance this approach to engineer the first targeted pan-RAS inhibitor for cancer therapy.

**One Sentence Summary:** Engineered chimeric toxin halts tumor growth *in vivo* via RAS cleavage

## Introduction

More than one-third of all human cancers harbor activating mutations in *RAS* oncogenes. Among the major isoforms (*H*-, *N*-, and *KRAS*), *KRAS* is the most frequently mutated oncogene and is found in nearly 25% of human malignancies and 85% of all RAS-driven cancers (Cox and Der, 2010; Hobbs et al., 2016; Prior et al., 2012). Notably, three out of the four deadliest cancers (pancreas, colorectal and lung) exhibit a high frequency of *KRAS* driver mutations (Hobbs et al., 2016; Prior et al., 2012). Moreover, *NRAS* and *HRAS* are also known oncogenic drivers in other neoplasms (Hobbs et al., 2016). Activating point mutations in *RAS* genes (mostly occurring at codon 12, 13 and 61) impair the intrinsic capacity of RAS proteins to hydrolyze GTP, thus locking them in a constitutively-activated GTP-bound state resistant to GAP-stimulated GTP hydrolysis. This leads to constitutive activation of downstream transduction signaling networks, such as the RAF/MEK/ERK (MAPK) axis, that drive survival and uncontrolled proliferation (Bos et al., 2007; Downward, 2003; Schubbert et al., 2007). Even in the absence of gain-of-function mutations, *RAS* genes still play a major role in tumorigenesis due to hyperactivation of the RAS/MAPK pathway via overexpression of upstream receptor tyrosine kinases (RTKs) and/or amplification of wild-type *RAS*, such as in head and neck squamous cell carcinoma (Hoa et al., 2002), esophageal and gastric cancers (Dulak et al., 2012), ovarian adenocarcinoma (Ross et al., 2013) and triple-negative breast cancer (TNBC) (Cancer Genome Atlas, 2012; Eckert et al., 2004).

Due to their major role in oncogenesis and progression of a wide spectrum of cancers via mutation-dependent and -independent mechanisms, RAS proteins have become a primary target for drug discovery and extensive effort has been directed to the development of selective RAS inhibitors (Papke and Der, 2017). However, the extremely high affinity for GTP and the absence of drug-accessible binding pockets have complicated efforts to directly inhibit RAS for decades, earning RAS the moniker “undruggable” (Cox et al., 2014; McCormick, 2015; Papke and Der, 2017). Nevertheless, recent success in the field has been achieved by selective targeting of mutant KRAS G12C with small molecules that covalently bind to the Cys12 residue in the KRAS switch-II pocket (Janes et al., 2018; Ostrem et al., 2013; Patricelli et al., 2016), and clinical trials are underway to validate their effectiveness (Amgen, 2019; Janssen Research & Development, 2019; Mirati Therapeutics Inc., 2019). However, these pharmacophores are specific for the G12C mutant form of KRAS and cannot be expanded to other mutants. Further, *KRAS^G12C^* mutations account for only about 11% of all *KRAS* mutations in cancer (O’Bryan, 2019) and are mainly detected in LAC (14%), followed by CRC (5%) and PDAC (1-3%) (Cox et al., 2014; Janes et al., 2018). Therefore, there remains an urgent need for a broadly applicable pan-RAS inhibitor for use against all RAS-driven tumors, either mutation-dependent or -independent.

Recently, we discovered a <u>R</u>AS/<u>R</u>AP1 <u>s</u>pecific endo<u>p</u>eptidase (RRSP) from *Vibrio vulnificus* that site-specifically cleaves RAS and its close homolog RAP1 between residues Y32 and D33 within the Switch I, a region crucial to RAS-mediated signal transduction. RRSP is highly specific for RAS and RAP1 and does not cleave other closely related GTPases (Antic et al., 2015; Biancucci et al., 2018; Biancucci et al., 2017). Importantly, RRSP cleaves all three major RAS isoforms (H, N, and K), as well as oncogenic RAS with mutations at position 12, 13 and 61. RRSP also targets both active (GTP-bound) and inactive (GDP-bound) RAS, resulting in destruction of the entire cellular RAS pool. By proteolytically cleaving the Switch I loop, RRSP prevents RAS from undergoing GDP-GTP exchange and binding the downstream effector kinase RAF, ultimately terminating ERK signaling in cells (Antic et al., 2015; Biancucci et al., 2018).

Assessing the therapeutic potential of RRSP is however precluded by the fact that RRSP is a 56-kDa domain of a larger protein toxin that alone does not readily diffuse across biological membranes, thus preventing its access to intracellular RAS. Recently, we demonstrated that the translocation machinery of diphtheria toxin (DT) can be engineered to deliver a broad diversity of passenger proteins into target cells (Auger et al., 2015; Park et al., 2018). In the present study, we exploit the receptor-targeting and membrane translocation properties of DT to deliver RRSP into target cells. This novel use of a translocating toxin is, in principle, similar to recombinant immunotoxins, which consist of bacterial toxins that have been re-targeted to tumor cells. For instance, Tagraxofusp and Ontak are composed of DT residues 1-389 fused to IL-3 and IL-2, respectively. Mechanistically, these drugs bind specific receptor-bearing cells (IL-3R and IL-2R, in the above example) and rely on the membrane translocating ability of DT to deliver an ADP-ribosyltransferase (DT_A_ domain) that targets elongation factor 2, resulting in termination of protein synthesis and subsequent cell death. These immunotoxins are extremely potent and are indicated for treatment of blastic plasmacytoid dendritic-cell neoplasm (Tagraxofusp) and cutaneous T-cell lymphoma (Ontak) (Jen et al., 2019; Prince et al., 2010).

In this study, we aimed to advance the recombinant immunotoxin concept by altering the DT delivery machinery payload to address the critical need for therapies that target RAS. To this end, we created a novel engineered chimeric toxin that uses the DT translocation system to deliver an enzyme that potently and irreversibly destroys RAS. We demonstrate the anti-cancer properties of this engineered chimeric toxin both *in vitro* and *in vivo*, providing proof-of-concept for the therapeutic development of RRSP as the first ever pan-RAS inhibitor.

## Results

### Engineered diphtheria toxin binding and translocation domain DT_B_ efficiently delivers RRSP into cells

RRSP is a 56-kDa bacterial toxin effector domain that requires a translocation machinery to cross cell membranes. To deliver RRSP into cells for assessment of its antitumorigenic activity, we have used an innovative DT-based protein delivery system. DT consists of a catalytically-active A fragment (DT_A_) and a B fragment (DT_B_) that includes both a receptor binding domain (DTR) and a translocation domain (DTT). DT_B_ alone can bind its surface receptor, the heparin-binding epidermal growth factor-like growth factor (HB-EGF), and transfer a wide range of protein cargos to the cell cytosol via a receptor-mediated endocytic mechanism (Auger et al., 2015). The introduction of two amino acid substitutions (K51E and E148K) into DT_A_ resulted in a non-toxic, catalytically-inactive A-fragment, herein referred to as DTa. As a first strategy to examine RRSP intracellular delivery via the DT platform, we expressed RRSP fused at the amino terminus of DTa with an intervening (G_4_S)_2_ linker, a design we previously used to successfully deliver a variety of other protein cargo (Auger et al., 2015; Park et al., 2018). Western blotting with the RAS10 pan-RAS antibody showed that treatment with RRSP-(G_4_S)_2_-DTa-DTT-DTR resulted in depletion of RAS in HCT-116 (*KRAS^G13D^*) cells at 1 nM (Figure 1A and Figure S1A). Despite this success, an intrinsic feature of this first-generation design is that RRSP is co-delivered with the DTa domain. We hypothesized that co-delivery of RRSP with inactive DTa could affect the catalytic activity of RRSP, either sterically or by DTa binding to its cytosolic EF2 target.

**Figure 1.**
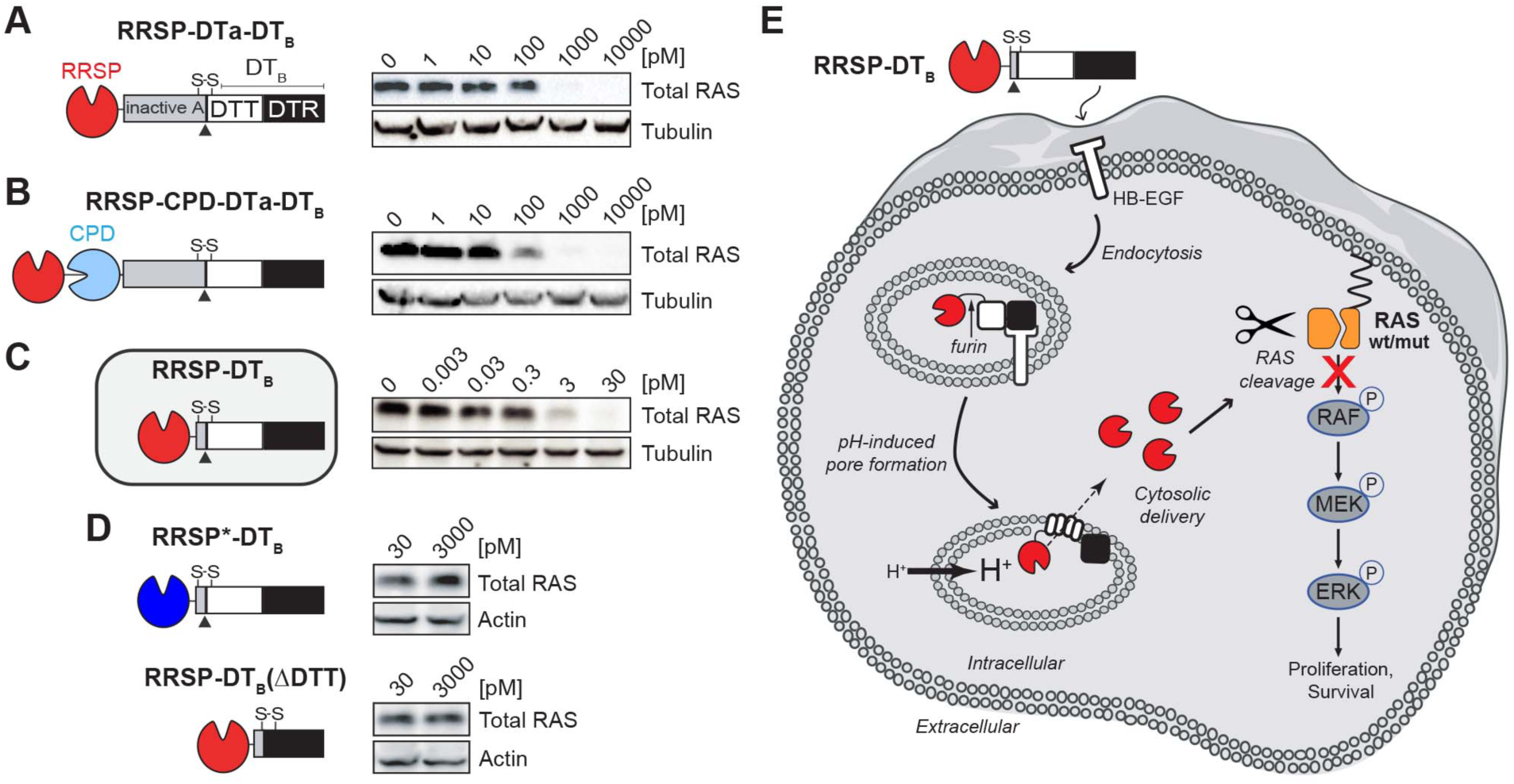
Engineered diphtheria toxin delivery machinery enables intracellular delivery of RRSP. Schematic of chimeric fusions of RRSP to (**A**) the amino terminus of non-toxic, full-length diphtheria toxin (RRSP-DTa-DTT-DTR), (**B**) to autoprocessing cysteine protease domain (CPD) of MARTX toxin from *Vibrio vulnificus* in the absence of the DTa domain (RRSP-CPD-DTT-DTR), and (**C**) to DTT-DTR without the DTa domain (RRSP-DTT-DTR or RRSP-DT_B_) and corresponding immunoblots against total RAS of lysates prepared from HCT-116 cells treated with RRSP-DT variants for 24 hours. (**D**) Immunoblots showing that the catalytically-dead RRSP*-DT_B_ mutant and the translocation-deficient control (RRSP-DTR) do not affect RAS levels in HCT-116 cells. (**E**) Schematic diagram illustrating transport and intracellular trafficking of RRSP fused to the translocation T and receptor R binding domains of diphtheria toxin fragment B (DT_B_) via a receptor-mediated endocytic mechanism. Once in the cytosol, RRSP cleaves RAS resulting in downregulation of the MAPK/ERK signaling cascade. Empty cell and vesicles were taken from Servier Medical Art database (https://smart.servier.com). HB-EGF, human heparin-binding epidermal growth factor-like growth factor.

To generate chimeras that would deliver RRSP alone into cells without DTa, we replaced DTa with an autoprocessing cysteine protease domain (CPD) from *Vibrio vulnificus* MARTX toxin that autocleaves and thereby releases its toxin effectors upon binding inositol hexakisphosphate in the host cytosol (Egerer and Satchell, 2010). When added to cells, RRSP-CPD-DTT-DTR improved RAS cleavage (partial inactivation at 100 pM) (Figure 1B and Figure S1B). In the second iteration design, we removed DTa, and appended RRSP via a (G_4_S)_2_ linker to the native release machinery of DT that includes DTT and DTR and named it RRSP-DT_B_. Indeed, cells treated with this much smaller protein showed complete loss of detectable RAS at concentrations as low as 3 pM (Figure 1C and Figure S1C). To confirm that RRSP-DT_B_ was working via the expected mechanism, we replaced RRSP with the catalytically-inactive mutant RRSP_H4030A_ (RRSP*-DT_B_) and we also deleted the translocation domain (DTT) required for cargo delivery (RRSP-DTR). Both control proteins failed to deplete RAS from cells (Figure 1D). The uptake of RRSP-DT_B_, translocation to cytosol, and subsequent RAS cleavage is summarized in Figure 1E. Together, these results demonstrate that DT_B_ is highly effective for intracellular translocation of RRSP across membranes to target RAS within cells. This final chimeric recombinant toxin was purified in large scale (Figure S2A-H), enabling the experimental investigation of the anti-cancer potential of RRSP.

### RAS processing by RRSP strongly affects viability and proliferation of TNBC cancer cells with activated wild-type RAS

We first investigated the anti-cancer potential of RRSP-DT_B_ in TNBC, a devastating disease and the last frontier in breast cancer research (Metzger-Filho et al., 2012). Activation of the RAS pathway remains an understudied field in breast cancer, most likely because of the low frequency of RAS mutations (∼5%) (Eckert et al., 2004). However, it has been shown that RAS proteins can be pathologically activated in TNBC via upregulation of RTKs and/or amplification of wild-type *RAS* that ultimately increase signaling through the RAS/MAPK pathway (Adeyinka et al., 2002; Eckert et al., 2004; Giltnane and Balko, 2014). The basal-like MDA-MB-436 cell line features overexpression of *KRAS*^WT^ and hyperactivation of ERK signaling (Mokhlis et al., 2019). MDA-MB-436 cells were treated with increasing concentrations of RRSP-DT_B_ for 1 and 24 hours and levels of intracellular RAS and phosphorylated ERK1/2 (pERK) were measured by western blot. Treatment with RRSP-DT_B_ resulted in complete cleavage of RAS at 10 nM after 1 hour, while 0.01 nM RRSP-DT_B_ was sufficient to completely cleave RAS after 24 hours. At all concentrations, levels of pERK were reduced in lockstep with RAS cleavage (Figure 2A). While pERK levels were markedly reduced after 24 hours, effects on cell viability reached a maximum at 72 hours, (IC_50_ = 0.005 ± 0.001 nM) (Figure 2B and S3A). In agreement with these results, the crystal violet staining assay showed cell loss starting at 0.01 nM after a 72-hour exposure to RRSP-DT_B_ (Figure 2C and Figure S3B), while a colony formation assay showed that RRSP-DT_B_ had a significant effect on cell proliferation at 0.01 nM and completely prevented colony formation at 10 nM (Figure 2D) over 10 days. Phenotypically, RRSP-DT_B_ caused significant cell rounding in MDA-MB-436 cells at 0.01 nM (Figure 2E). Therefore, RRSP-DT_B_ causes highly potent cytotoxic effects via the efficient ablation of RAS signaling in a *KRAS^WT^* cell line.

**Figure 2.**
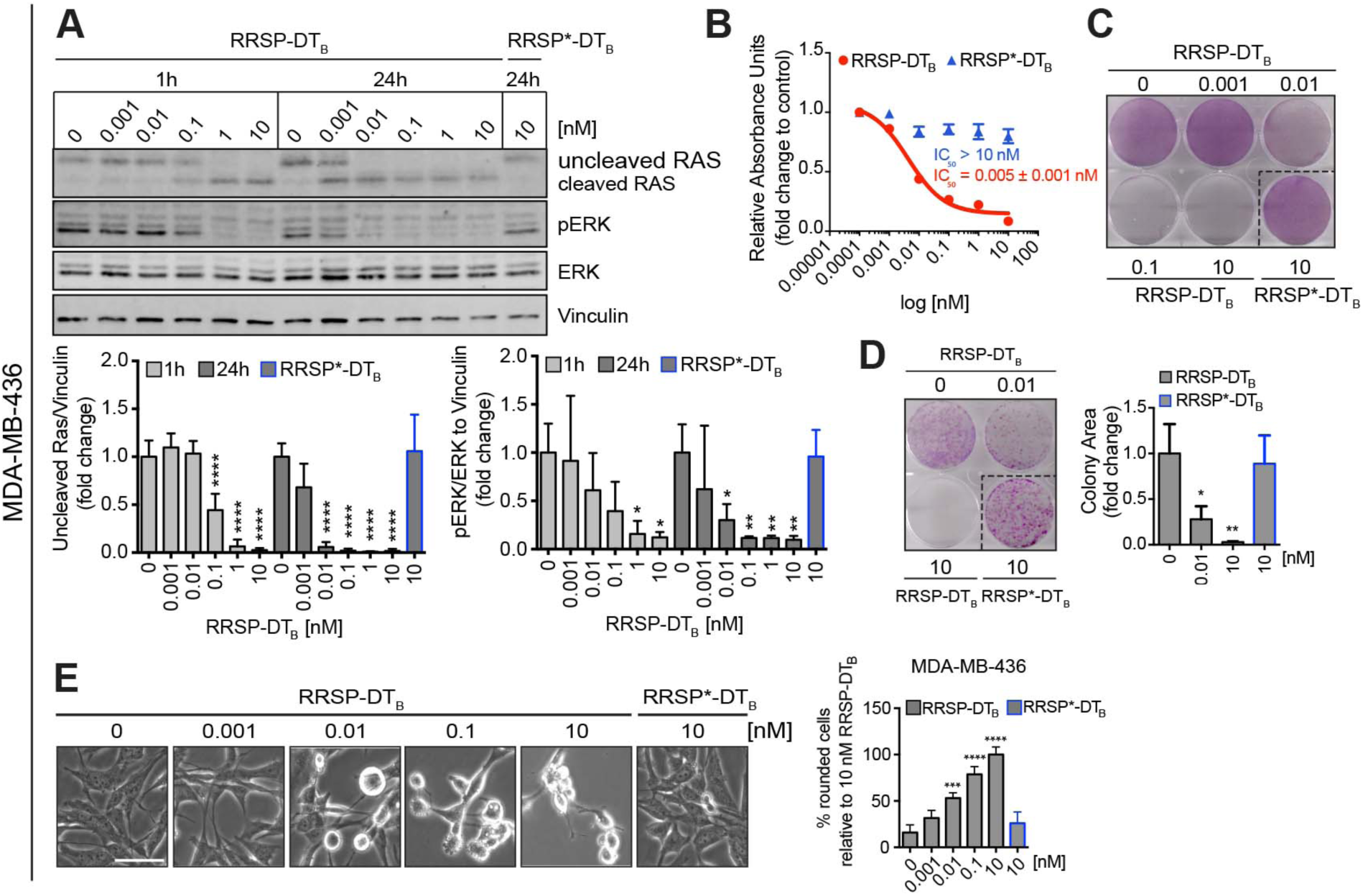
RAS processing by RRSP leads to reduced viability and proliferation of a triple-negative *KRAS^WT^* breast cancer cell line. (**A**). Representative western blot and densitometric analysis of uncleaved RAS and phosphorylated ERK (hereafter pERK) levels in MDA-MB-436 *KRAS^WT^* cells treated with increasing concentrations of RRSP-DT_B_ for 1 and 24 hours. The catalytically-inactive RRSP*-DT_B_ mutant was used as negative control at 10 nM and vinculin as gel loading control. Results are expressed as means ± SD of four independent experiments (\**p* < 0.05, \*\**p* < 0.01, \*\*\**p* < 0.001, \*\*\*\**p* < 0.0001 versus corresponding control 0 nM; one-way ANOVA followed by Dunnett’s multiple comparison test, *n*= 4). (**B**). Fitted dose-response curve of RRSP-DT_B_ in MDA-MB-436 cells following 72 hours of treatment. Results are expressed as mean ± SEM (*n*=3). (**C**). Representative images of crystal violet staining of MDA-MB-436 cells treated with RRSP-DT_B_ or RRSP*-DT_B_ as indicated for 72 hours. (**D**). Representative images and quantitative analysis of crystal violet-stained colonies from MDA-MB-436 cells pre-treated with RRSP-DT_B_ and RRSP*-DT_B_ at the indicated concentrations for 72 hours and replated at 2,500 cells/well to form colonies over 10 days. Results are expressed as means ± SD of three independent experiments (\**p* < 0.05, \*\**p* < 0.01 versus corresponding control 0 nM; one-way ANOVA followed by Dunnett’s multiple comparison test, *n*= 3). (**E**) Bright field images of MDA-MB-436 cells treated with RRSP-DT_B_ and RRSP*-DT_B_ at the indicated concentrations for 48 hours (scale bar = 50 µM). Up to 200 rounded cells (with sharp edges and no protrusions) per image were counted manually and means ± SD of three independent experiments were plotted. Because all cells treated with 10 nM of RRSP-DT_B_ appeared round, this condition was set as 100% cell roundness (\*\*\**p* < 0.001, \*\*\*\**p* < 0.0001 versus corresponding control 0 nM; one-way ANOVA followed by Dunnett’s multiple comparison test, *n*= 3).

### RRSP halts tumor growth of a *KRAS^WT^* TBNC xenograft

As RRSP-DT_B_ exhibited a potent effect on the viability of MDA-MB-436 cells *in vitro*, we next investigated how it affected tumor growth *in vivo*. Retaining the native receptor targeting of DT_B_ to translocate RRSP is advantageous as a test system for mouse xenografts since DT is at least 1000-fold less potent on cells expressing murine HB-EGF than cells expressing human HB-EGF (Cha et al., 2003; Mitamura et al., 1995; Palmiter, 2001). Indeed, treatment with RRSP-DT_B_ did not affect the viability of mouse embryonic fibroblasts (MEFs) (Figure S4A-B). Thus, DT_B_ in the chimeric toxin functions as a cancer cell targeting domain in xenograft models since human HB-EGF is expressed only on the xenografted human cells (Figure 3A). A maximum tolerated dose (MTD) study determined that 0.1 mg/kg administered intraperitoneally (IP) every day (q.d.) was well tolerated, while dosage at or above 0.5 mg/kg caused weight loss (Figure S4C-D). In the MDA-MB-436 xenograft, IP administration of RRSP-DT_B_ at 0.1 mg/kg every other day (weekends excluded) for 4 weeks halted tumor growth (Figure 3A and 3B). Importantly, three out of five mice treated with RRSP-DT_B_ showed tumor regression by the end of the treatment schedule (Figure 3C). Most importantly, catalytically-inactive RRSP*-DT_B_ did not affect tumor growth compared to the vehicle (saline) control, demonstrating that tumor regression was due specifically to the RAS processing activity of RRSP. Indeed, RRSP-DT_B_ dramatically reduced total RAS immunoreactivity in residual tumors, confirming effective RAS cleavage *in vivo* (Figure 3D-E and Figure S5A). Additional experimental groups using slight variations of the dosing strategy corroborated these findings (Figure S6A-F). Since ERK phosphorylation was limited to the outer rim of control MDA-MB-436 tumors excised from mice at 4 weeks (Figure S5B and Figure S6G), we assessed pERK levels at an earlier time point when the smaller size of large control tumors is more amenable to pERK detection. In tumors harvested at two weeks, we observed staining throughout the section with high basal pERK levels in both saline and RRSP*-DT_B_-treated tumors and with significant reduction in pERK in the RRSP-DT_B_ treatment group (Fig S5C). These results demonstrate that RRSP-DT_B_ effectively engages and ablates RAS *in vivo*, resulting in decreased ERK activation and tumor regression.

**Figure 3.**
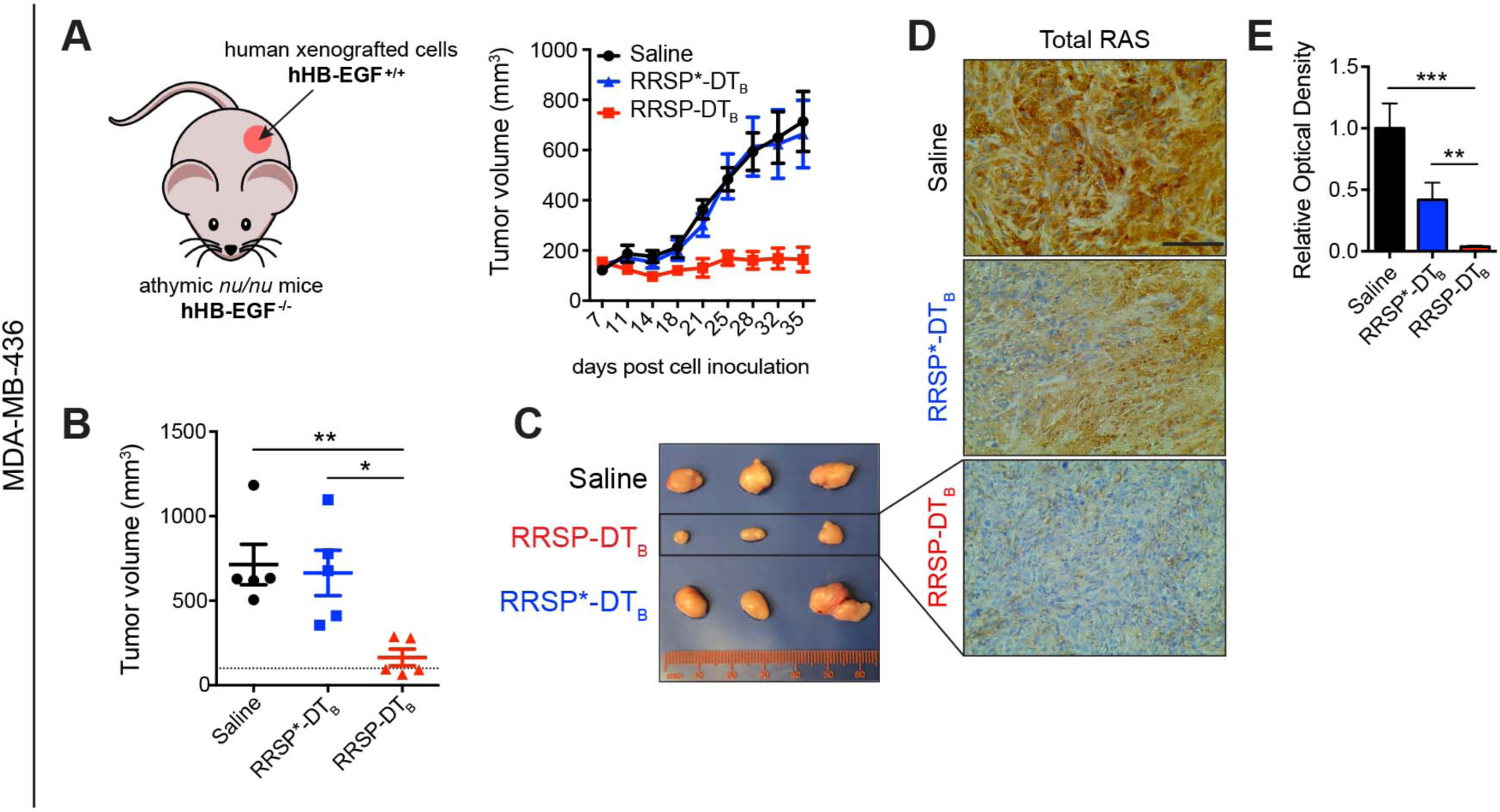
RRSP halts tumor growth in a *KRAS^WT^* TNBC xenograft *in vivo*. (**A)**. Tumor growth curve of vehicle, RRSP-DT_B_ and RRSP*-DT_B_-treated athymic *nu/nu* female mice bearing MDA-MB-436-derived tumors. Mice received 0.1 mg/kg of RRSP-DT_B_ and 0.1 mg/kg of RRSP*-DT_B_ every other day (weekends excluded) via IP injection. (**B**). Representative images of MDA-MB-436 tumors at the experimental endpoint. (**C**). Column scatter plots showing individual tumor volumes at the end of the treatment schedule for MDA-MB-436 xenografts. Horizontal dashed line indicates the baseline tumor volume on the first day of treatment, which corresponds to the average of tumor volumes at the indicated time (140 mm^3^). Data are means ± SEM (*n*=5 mice per group; \**p* < 0.05, \*\**p* < 0.01). (**D**). Representative images of immunoreactivity to total RAS in sections from MDA-MB-436 tumors and (**E**) corresponding quantification of DAB optical density via color deconvolution using whole tumor sections (\*\**p* < 0.01, \*\*\**p* < 0.001, one-way ANOVA followed by Tukey’s multiple comparison test, *n*=3; scale bar = 100 µM).

### Assessment of relative susceptibility to RRSP-DT_B_ using the NCI-60 panel

Following the positive results obtained with the wild-type *RAS* MDA-MB-436 cell line both *in vitro* and *in vivo*, we screened RRSP-DT_B_ on the National Cancer Institute NCI-60 human tumor cell line panel comprised of 60 cell lines representing nine different cancer types with various genetic backgrounds, including *RAS* mutations.

Growth inhibition caused by RRSP-DT_B_ at the highest dose employed (13.5 nM) was measured by sulforhodamine B assay after 48 hours. All data were normalized such that 0% growth inhibition corresponded to no change in the cell number relative to control after 48 hours while a drop of 100% corresponded to no cell growth relative to time zero (100% growth inhibition); values below 100% correspond to cell loss. Fourteen cell lines were classified as “highly susceptible” to RRSP-DT_B_ as they showed growth inhibition greater than or equal to 90%. Thirty-eight cell lines showed varying degrees of growth inhibition from 25 to 90% and were designated as “susceptible” to RRSP-DT_B_ (Figure 4A).

**Figure 4.**
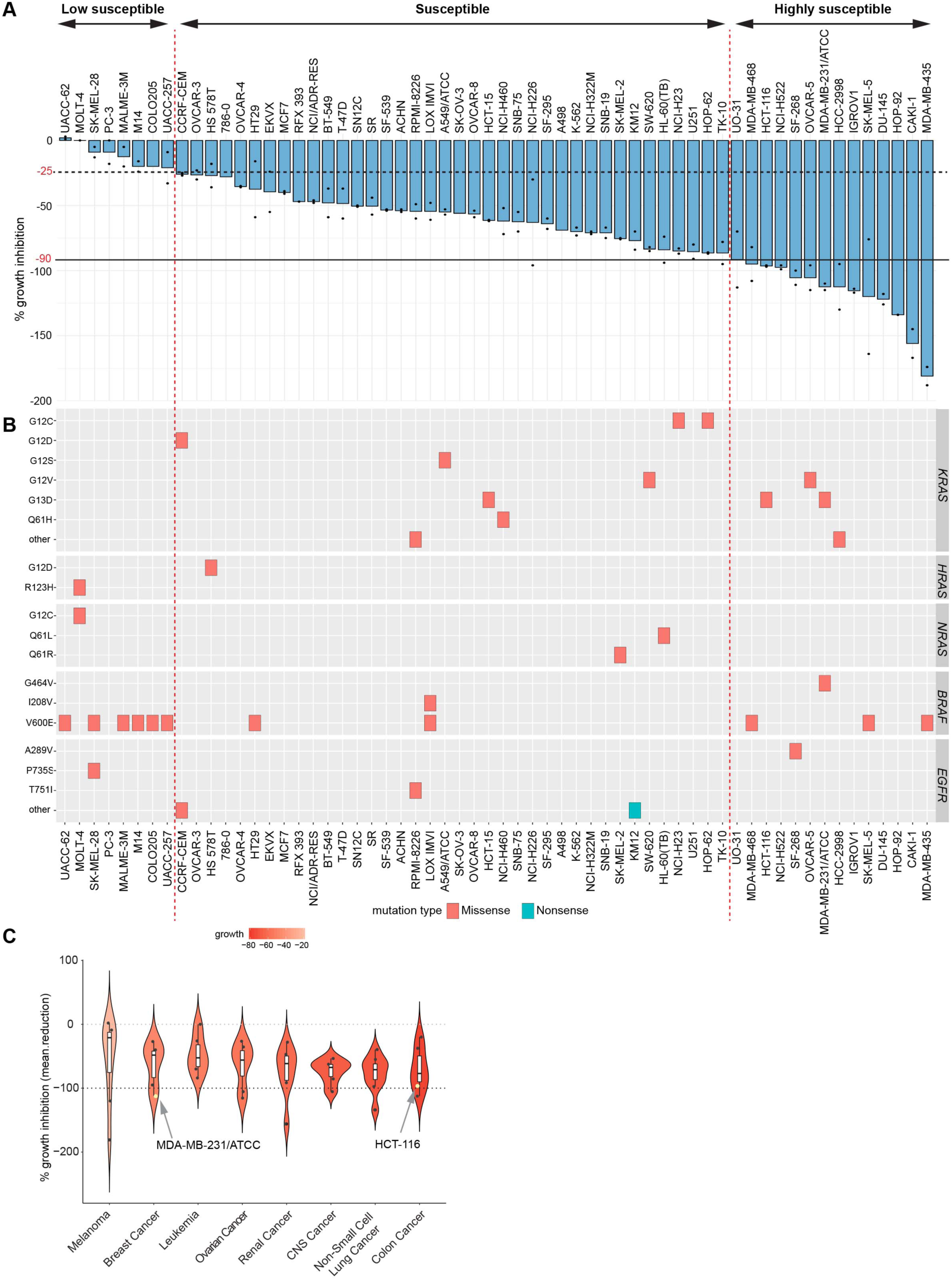
Comprehensive analysis of the effect of RRSP on the NCI-60 cancer cell line panel. (**A**). Bar plot showing the percent growth inhibition of RRSP-DT_B_ on the NCI-60 panel. Cancer cell lines were ranked in descending order based on their growth inhibition % value. The presence of two dots on the bars indicate that two replicates were performed per each cell line and bars represent means. No dots indicate that only one replicate was available. (**B**). Spectrum of missense and nonsense mutations in *KRAS, HRAS, NRAS, BRAF* and *EGFR* genes in the 60 cell lines of the panel. (**C**). Violin plot showing the median percent growth inhibition of cell lines treated with RRSP-DT_B_ and grouped per tumor type in descending order. Only tumor types that had at least 5 cell lines per group were plotted in the graph.

Generally, cell lines from the panel that carry *KRAS* missense mutations were among the most responsive to RRSP-DT_B_, while cell lines with mutations in *BRAF* (especially *BRAF^V600E^*) tended to be less responsive (Figure 4B). Mutations in *HRAS*, *NRAS,* or *EGFR* were not associated with a response pattern to RRSP-DT_B_ treatment (Figure 4B). Notably, analysis of copy number alterations from exome data available for 53 out of 60 NCI cell lines showed amplification of *KRAS* and *NRAS,* as well as deletions in *EGFR*, in the cell lines most sensitive to RRSP-DT_B_ (Figure S7B-C). Further, colon cancer cell lines as a group were the most sensitive to RRSP-DT_B_overall, followed by non-small cell lung cancer lines (Figure 4C).

Eight cell lines in the NCI-60 screen showed <25% growth inhibition compared to mock treated and were categorized as “less susceptible”. In this group, growth of UACC-62 and MOLT-4 cells was not affected by RRSP-DT_B_. One requirement for RRSP-DT_B_ cytotoxicity is expression of the DT receptor HB-EGF on the cell surface. Analysis of available *HBEGF* gene expression data (Figure S7A) showed that, while the correlation between RRSP-DT_B_ sensitivity and *HBEGF* expression is not linear, many of the cell lines that responded to RRSP-DT_B_ had higher expression of *HBEGF*. In addition, the TNBC Hs578T cell line *(HRAS^G12D^)*, categorized as less sensitive in the NCI-60 screen, was confirmed to have lower expression of HB-EGF protein and a moderate reduction in cell viability after 72 hours, although RAS cleavage and ERK dephosphorylation were detected at earlier time points (Figure S8A-E). Thus, when taken altogether, results from the NCI-60 screen indicate that most tumor types showed sensitivity to RRSP-DT_B_ and cell lines that display genomic abnormalities in *RAS* genes were markedly sensitive to RRSP-DT_B_ treatment.

### RRSP reduces tumor burden in a mutant *KRAS* TNBC xenograft model

The highly sensitive basal-like MDA-MB-231 TBNC cell line *(KRAS^G13D^)* in the NCI-60 screen (Figure 4A) is a *KRAS*-dependent cell line (Scholl et al., 2009), displays high KRAS protein expression levels (Mokhlis et al., 2019), and was among the first cell lines characterized as sensitive to RRSP (Antic et al., 2015, Figure 4A). Consistent with these findings, treatment of MDA-MB-231 cells with the engineered chimeric toxin RRSP-DT_B_ cleaved RAS and reduced pERK levels to a similar extent as in MDA-MB-436 cells (Figure 5A). Also similar to MDA-MB-436 cells, cell viability (Figure 5B-C and Figure S9A-B) and cell proliferation (Figure 5D) were strongly affected after 72 hours (IC_50_ = 0.012 ± 0.001 nM), and RRSP-DT_B_ induced significant cell rounding in MDA-MB-231 cells starting at 0.1 nM (Figure 5E). To further investigate these results, we tested RRSP-DT_B_ in a xenograft study using the TNBC MDA-MB-231 *(KRAS^G13D^)* cell line. This cell line has a more rapid doubling time than MDA-MB-436 cells (Hassan et al., 2017), so mice were treated with a more frequent 5 days ON/2 days OFF schedule for 4 weeks, resulting in significant anti-tumor activity (Figure 5F). After 4 weeks of treatment with 0.1 mg/kg q.d. of RRSP-DT_B_, the treatment group had markedly smaller tumors than saline and RRSP*-DT_B_ controls (Figure 5G-H). RAS and pERK immunostaining showed that RRSP-DT_B_-treated tumors had a more focal/patchier staining pattern than tumors from saline and RRSP*-DT_B_ treated mice, which instead exhibited a more diffuse staining pattern (Figure S9C-D). Indeed, MDA-MB-231 tumors (Figure 5G) were pale in color compared to the MDA-MB-436 tumors (Figure 3B) consistent with reports that MDA-MB-231 tumors exhibit low vascularization (Fleming et al., 2010). This supports that MDA-MB-231 tumors in our study may be poorly vascularized, and tumor size reduction may be predominantly due to RRSP-DT_B_ diffusion from the tumor periphery, resulting in the partial cleavage of RAS in the center of residual tumors. Even still, RRSP-DT_B_ was highly effective in targeting TBNC MB-MDA-231 xenografts resulting in a significant inhibition of tumor growth and confirmed that RRSP-DT_B_ is effective against TNBC.

**Figure 5.**
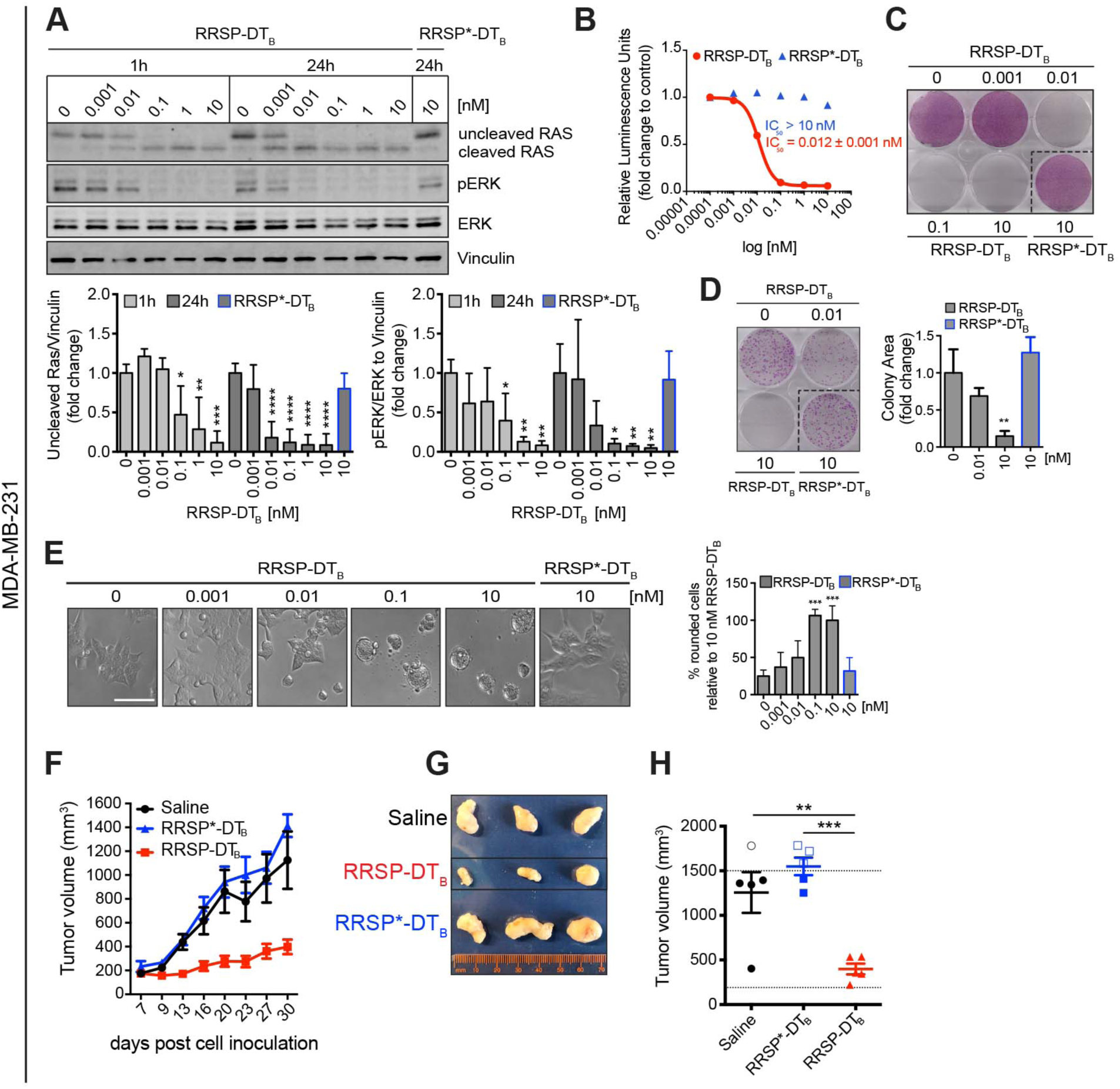
RRSP inhibits cell viability, proliferation and tumor growth in a *KRAS^G13D^*-dependent TNBC xenograft *in vivo.* (**A**). Representative western blot and densitometric analysis of uncleaved RAS and pERK levels in MDA-MB-231 *KRAS^G13D^* cells treated with increasing doses of RRSP-DT_B_ and with 10 nM of the catalytically-inactive RRSP*-DT_B_. Results are expressed as means ± SD of four independent experiments (\**p* < 0.05, \*\**p* < 0.01, \*\*\**p* < 0.001, \*\*\*\**p* < 0.0001 versus corresponding control 0 nM; one-way ANOVA followed by Dunnett’s multiple comparison test, *n*= 4). (**B**). Fitted dose-response curve of RRSP-DT_B_ in MDA-MB-231 cells following 72 hours of treatment. Results are expressed as means ± SEM (*n*=3). (**C**). Representative images of crystal violet staining of MDA-MB-231 cells treated with RRSP-DT_B_ or RRSP*-DT_B_ as indicated for 72 hours. (**D**). Representative images and quantitative analysis of crystal violet-stained colonies from MDA-MB-231 cells pre-treated with RRSP-DT_B_ and RRSP*-DT_B_ at the indicated concentrations for 72 hours and replated at 2,500 cells/well to form colonies over 10 days. Results are expressed as means ± SD of three independent experiments (\*\**p* < 0.01 versus corresponding control 0 nM; one-way ANOVA followed by Dunnett’s multiple comparison test, *n*= 3). (**E**) Bright field images of MDA-MB-231 cells treated with RRSP-DT_B_ and RRSP*-DT_B_ at the indicated concentrations for 48 hours (scale bar = 50 µM). Up to 200 rounded cells per image were counted manually and means ± SD of three independent experiments were plotted (\*\*\**p* < 0.001 versus corresponding control 0 nM; one-way ANOVA followed by Dunnett’s multiple comparison test, *n*= 3). (**F**). Tumor growth curve of vehicle, RRSP-DT_B_ and RRSP*-DT_B_-treated athymic *nu/nu* female mice bearing MDA-MB-231-derived tumors. Mice received 0.1 mg/kg of RRSP-DT_B_ and 0.1 mg/kg of RRSP*-DT_B_ every day (weekends excluded). (**G**). Representative images of MDA-MB-231 tumors at the experimental endpoint. (**H**). Column scatter plots showing individual tumor volumes at the end of the treatment schedule. Horizontal dashed line indicates the baseline tumor volume on the first day of treatment, which corresponds to the average of tumor volumes at the indicated time (194 mm^3^). Empty points indicate tumors from mice that were sacrificed earlier because they exceeded the 1500 mm^3^ threshold. Their size was recorded before euthanasia. Data are means ± SEM (*n*=5 mice per group). Statistical analysis between vehicle and treatment groups was performed using one-way ANOVA followed by Tukey’s multiple comparison test (\*\**p* < 0.01, \*\*\**p* < 0.001).

### RRSP-DT_B_ inhibits cell viability of colorectal cancer cells in 2D monolayers and 3D spheroids

Mutations in *KRAS* are detected in approximately 50% of patients with colorectal carcinoma (CRC), which is the third leading cause of cancer-related death in both men and women in the United States and represents a major target population for anti-RAS therapy (Ferlay et al., 2015). Interestingly, we found that cell lines from colon cancer were the most susceptible to RRSP-DT_B_ in the NCI-60 screen. Here, we examined the anti-cancer potential of RRSP-DT_B_ on the CRC cell line HCT-116 harboring a *KRAS^G13D^*mutation. Treatment of cells with RRSP-DT_B_ at 10 pM led to RAS processing and reduced pERK levels at 10 pM after 24 hours (Figure 6A) and strongly decreased the viability of HCT-116 cells after 72 hours (IC_50_ = 0.0015 ± 0.002 nM) (Figure 6B and S10A). Crystal violet staining showed remarkable cell loss after treatment with RRSP-DT_B_ as low as 10 pM (Figure 6C and S10B). Clonogenic assays showed a significant reduction in HCT-116 colonies following treatment with 10 pM of RRSP-DT_B_ and almost no colonies were found at 10 nM RRSP-DT_B_ (Figure 6D) demonstrating complete loss of cell proliferation. These data agree with the RRSP-DT_B_-induced cell rounding phenotype observed with this and other sensitive cell lines (Figure 6E).

**Figure 6.**
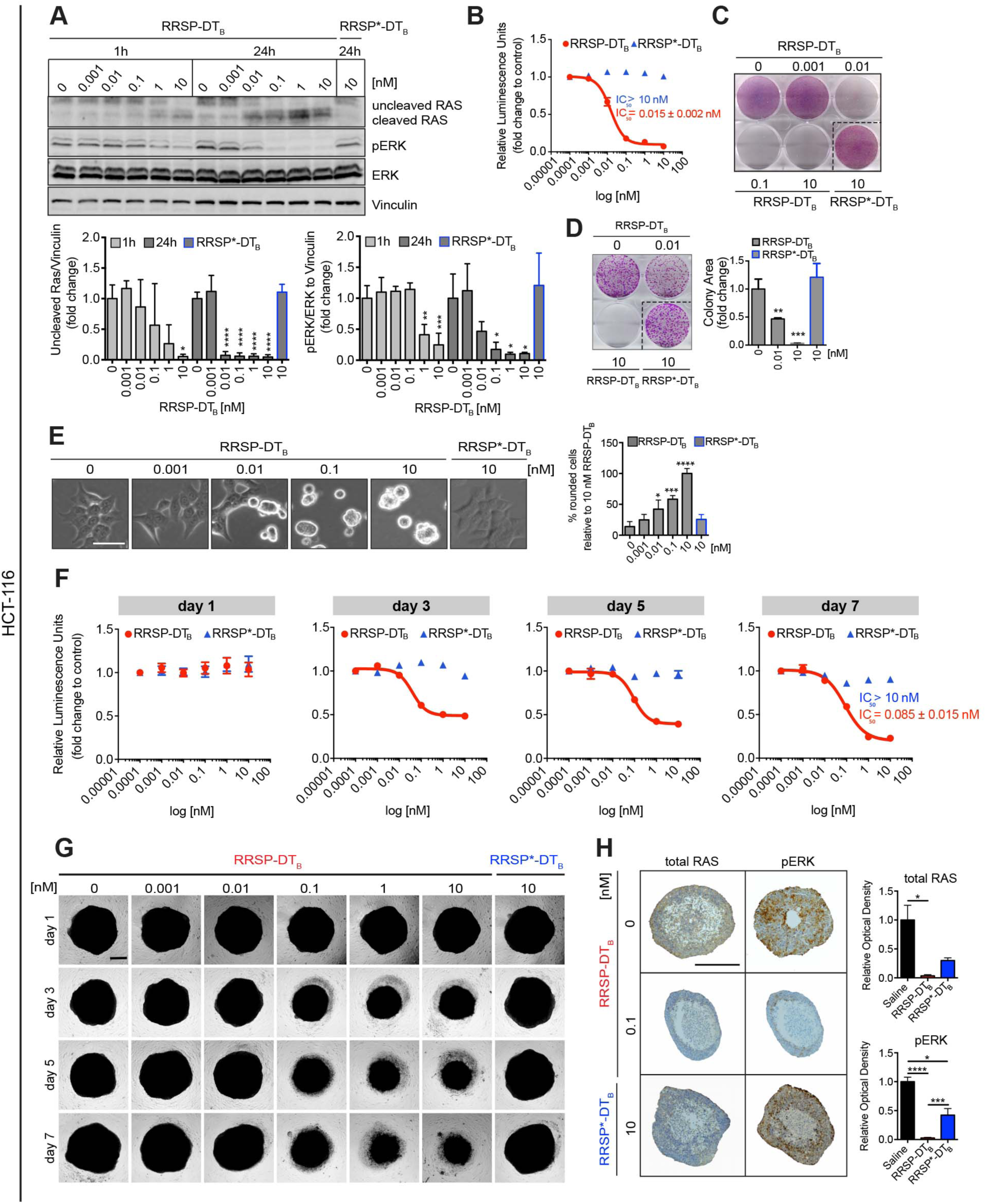
Effect of RRSP on viability and proliferation of colorectal HCT-116 *KRAS^G13D^* cells in 2D and 3D cellular models. (**A**) Representative western blot and densitometric analysis of uncleaved RAS and phosphorylated ERK in HCT-116 cells treated with increasing doses of RRSP-DT_B_ for 1 and 24 h. The catalytically-inactive RRSP*-DT_B_ mutant was used as negative control at 10 nM. Results are expressed as means ± SD of three independent experiments (\**p* < 0.05, \*\**p* < 0.01, \*\*\**p* < 0.001, \*\*\*\**p* < 0.0001 versus corresponding control 0 nM; one-way ANOVA followed by Dunnett’s multiple comparison test, *n* = 3). (**B**) Fitted dose-response curve of RRSP-DT_B_ in HCT-116 cells following 72 h of treatment. (**C**). Representative images of crystal violet staining of HCT-116 cells treated with RRSP-DT_B_ or RRSP*-DT_B_ as indicated for 72 h. (**D**) Representative images and quantitative analysis of crystal violet-stained colonies from HCT-116 cells pre-treated with RRSP-DT_B_ or RRSP*-DT_B_ at the indicated concentrations for 72 h and reseeded at 2,500 cells/well to form colonies over 10 days. Results are expressed as means ± SD of three independent experiments (\*\**p* < 0.01, \*\*\**p* < 0.001 versus corresponding control 0 nM; one-way ANOVA followed by Dunnett’s multiple comparison test, *n* = 3). (**E**). Bright field images of HCT-116 cells treated with RRSP-DT_B_ and RRSP*-DT_B_ at the indicated concentrations for 48 h. Up to 200 rounded cells per image were counted manually and means ± SD of three independent experiments were plotted. (\**p* < 0.05, \*\*\**p* < 0.001, \*\*\*\**p* < 0.0001 versus corresponding control 0 nM; one-way ANOVA followed by Dunnett’s multiple comparison test, *n* = 3). (**F**). Fitted dose-response curves in HCT-116 spheroids following treatment with RRSP-DT_B_ and RRSP*-DT_B_ at concentrated indicate for time indicate (d, day). Results are expressed as means ± SEM (*n*=4). (**G**) Representative bright field images of HCT-116 spheroids treated at the indicated time and concentrations with RRSP-DT_B_ and RRSP*-DT_B_ and quantitative analysis of spheroids’ volume (scale bar = 200 µm). Results are expressed as means ± SD of three independent experiments (\*\**p* < 0.01, \*\*\*\**p* < 0.0001 versus corresponding control 0 nM; one-way ANOVA followed by Dunnett’s multiple comparison test, *n* = 3). (**H**). Representative images of immunoreactivity to total RAS and phosphorylated ERK in sections from HCT-116 spheroids and corresponding quantification of DAB optical density via color deconvolution (\**p* < 0.05, \*\*\**p* < 0.001, \*\*\*\**p* < 0.0001; one-way ANOVA followed by Tukey’s multiple comparison test, *n*=3; scale bar = 400 µM).

Unlike two-dimensional cell monolayers, it has been shown that three-dimensional spheroids can recapitulate several architectural, microenvironmental and functional features of *in vivo* tumors while retaining reproducibility and easy-to-use properties (Ferreira et al., 2018; Vidimar et al., 2018). We generated spheroids from HCT-116 cells in ultra-low attachment multiwell plates and treated them with RRSP-DT_B_ for up to 7 days. Effects on cell viability were observed after 3 days, while the maximum effect was observed after 7 days of treatment (IC_50_ = 0.085 ± 0.015 nM) (Figure 6F). Bright field images of HCT-116 spheroids and quantitative analysis of their volume showed a dose- and time-dependent reduction in size following RRSP-DT_B_ treatment (Figure 6G and S10C). Immunostaining on sections of HCT-116 spheroids treated with RRSP-DT_B_ with 0.1 nM for 3 days (when spheroids began to regress) showed that intracellular RAS was essentially absent and pERK was undetectable (Figure 6H). Collectively, these results demonstrate that RRSP-DT_B_-dependent RAS ablation and subsequent loss of phosphorylated ERK is highly cytotoxic to CRC HCT-116 (*KRAS^G13D^)* cells in 2D monolayers and 3D spheroids.

### RRSP-DT_B_ exhibits antitumor activity in a colorectal cancer xenograft model

Since HCT-116 cells are fast-growing cells (doubling time ≤ 20 hours) that generate fast-growing tumors *in vivo* (Ahmed et al., 2013), we chose to administer 0.1 mg/kg of RRSP-DT_B_ to mice on a q.d. (1X/day) and twice per day (2X/day, b.i.d.) schedule (weekends excluded) for 4 weeks. Both dosing schedules resulted in a significant reduction in tumor size, while tumor regression was observed in 2/10 mice in both RRSP-DT_B_ treatment groups (Figure 7A-B). Analysis of residual tumors showed that RRSP-DT_B_ effectively cleaved RAS, although only the b.i.d. dosing group achieved statistical significance (Figure 7C-D and S11A). Quantitative analysis of pERK showed that the b.i.d. dosing group had a 2.5-fold reduction in pERK levels relative to controls and some tumors displayed focal staining patterns (Figure S11B). Overall, these data show that RRSP-DT_B_ exhibited strong anti-tumor activity in a CRC xenograft model via irreversible inactivation of RAS.

**Figure 7.**
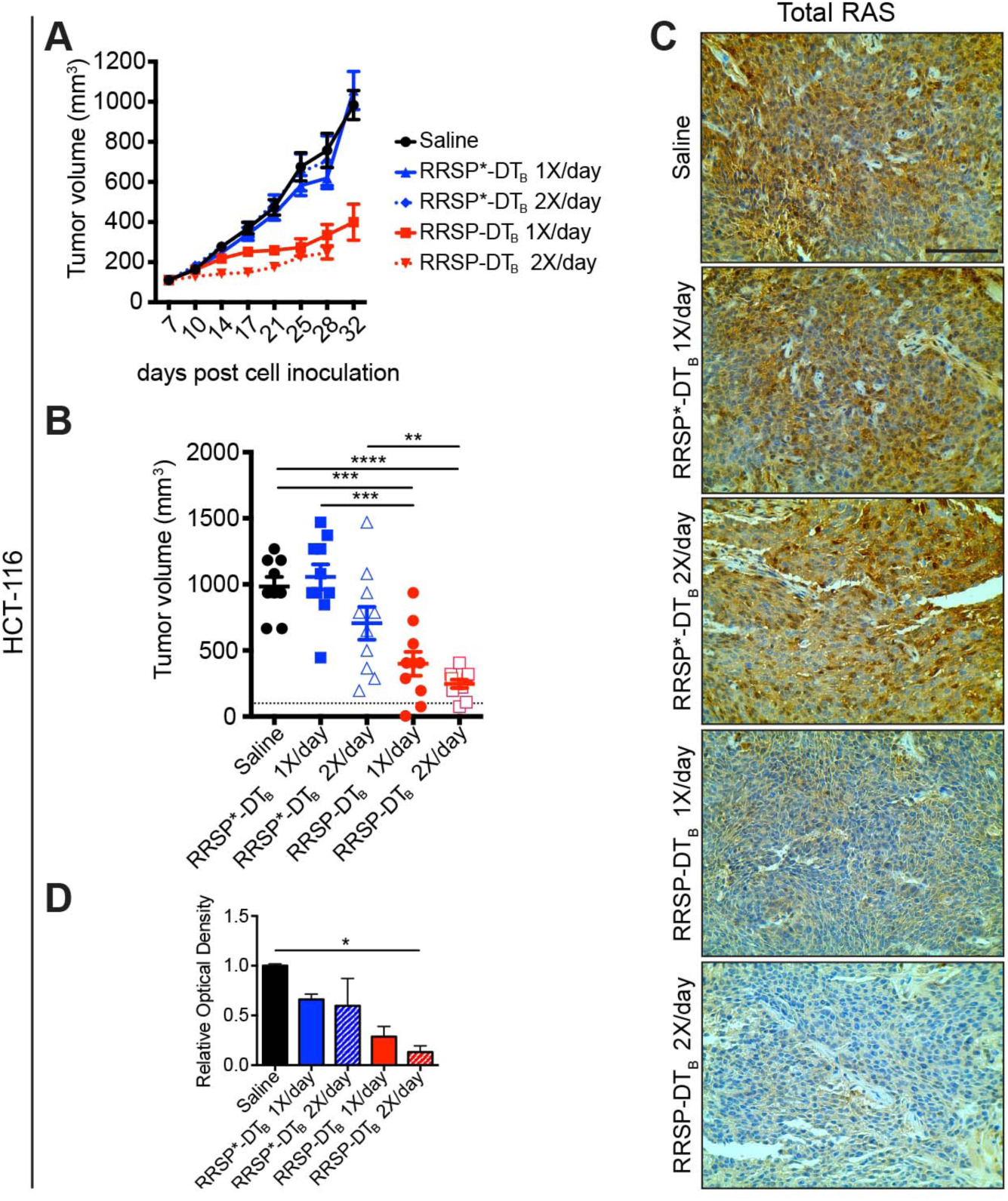
RRSP slows tumor growth in a CRC xenograft *in vivo*. (**A**) Tumor growth curve of vehicle, RRSP-DT_B_ and RRSP*-DT_B_-treated athymic *nu/nu* female mice bearing HCT-116-derived tumors. Mice received 0.1 mg/kg of RRSP-DT_B_ and 0.1 mg/kg of RRSP*-DT_B_ every day (1X/day) or twice per day (2X/day) weekends excluded. (**B**) Column scatter plots showing HCT-116 individual tumor volumes at the end of the treatment schedule. Horizontal dashed lines indicate the baseline tumor volume on the first day of treatment, which corresponds to 100 mm^3^. Data are means ± SEM. In all groups, *n*=10 mice. In the saline group, one mouse was sacrificed on day 28 due to a too large tumor, whose size was recorded before euthanasia. Statistical analysis between vehicle and treatment groups was performed using one-way ANOVA followed by Tukey’s multiple comparison test (\*\**p* < 0.01, \*\*\**p* < 0.001, \*\*\*\**p* < 0.0001). (**C**). Representative IHC images of immunoreactivity to total RAS in sections from HCT-116 tumors and (**D**) corresponding quantification of DAB optical density via color deconvolution using whole tumor sections (\**p* < 0.05, one-way ANOVA followed by Tukey’s multiple comparison test, *n*=3; scale bar = 100 µM).

## Discussion

Almost four decades ago, the discovery of RAS as the first human oncogene changed our understanding of cancer. Despite tremendous effort, the three RAS isoforms (KRAS, NRAS and HRAS) have been called “undruggable” and no direct therapies are currently in clinical use (Cox and Der, 2010; Cox et al., 2014; McCormick, 2015; Papke and Der, 2017). Nevertheless, promising results have been recently reported for small molecules that irreversibly bind the G12C mutant form of KRAS and phase I clinical trials are ongoing to evaluate the efficacy and safety profile of AMG510, MRTX849 and ARS3248 (Amgen, 2019; Canon et al., 2019; Hallin et al., 2019; Janssen Research & Development, 2019; Mirati Therapeutics Inc., 2019). However, the *KRAS^G12C^* mutation is found in only a subpopulation of cancers, limiting the applicability of these compounds. Further, while RAS oncoproteins remain the main oncogenic drivers in RAS-addicted tumors, several studies have pinpointed the tumorigenic role of wild-type RAS proteins (Zhou et al., 2016). Indeed, amplification of wild-type *RAS* genes or activation of wild-type RAS proteins via acute growth factor stimulation have been shown to sustain growth of multiple tumor types (Dulak et al., 2012; Eckert et al., 2004; Hoa et al., 2002; Ross et al., 2013). Moreover, it has been previously reported that depletion of mutant *RAS* in cancer cells harboring a heterozygous *RAS (K-, N-* or *H)* mutation leads to overactivation of EGFR/RAS signaling due to the remaining wild-type RAS isoforms (Young et al., 2013). Therefore, there is an urgent need for broadly applicable pan-RAS inhibitors that can target not only the most common RAS mutants, but also wild-type RAS proteins aberrantly overactivated by mutation-independent mechanisms.

Here, we describe a novel engineered chimeric toxin comprised of an endopeptidase from *Vibrio vulnificus* that is highly specific for RAS and RAP1 and the protein translocation machinery of diphtheria toxin. This fusion protein is able to mediate the endocytosis and cytosolic delivery of RRSP exclusively into receptor-bearing cells. We demonstrate that RRSP delivery by DT_B_ into HB-EGF-expressing cells results in cleavage of RAS in the picomolar range. As RRSP processes all forms of RAS, the entire intracellular pool of RAS is affected leading to dramatic loss of pERK and cell proliferation. RRSP-DT_B_ affected cell proliferation of many cancer cell lines that express HB-EGF on the cell surface. Of note, HB-EGF is highly expressed in several human cancers, including gastric, ovarian and triple-negative breast cancer, and the non-toxic diphtheria toxoid CRM197 had anti-tumor effects on TNBC xenografts (Nam et al., 2016). As such, we predicted TBNC could be highly susceptible to RRSP-DT_B_. We show that picomolar amounts of RRSP-DT_B_ completely cleaved RAS proteins, thus eliminating RAS-dependent ERK phosphorylation in these cells. This resulted in a very potent effect on cell viability (IC_50_ = 5 pM). Similarly, RRSP-DT_B_treatment led to a large reduction in tumor burden in MDA-MB-436 xenograft experiments, with all mice showing significantly smaller tumors than control groups and 60% of mice showing tumor regression. Target engagement was confirmed by immunohistochemistry analysis of tumor sections showing complete depletion of RAS in residual tumors. We also observed low pERK staining in small dissected RRSP-DT_B_-treated tumors, although pERK levels in these specimens were not significantly different than control tumors that showed low detection of pERK after four weeks. However, assessment of resected tumors from a shorter 14-day xenograft study revealed that RRSP-DT_B_ significantly reduced levels of pERK. This effect was masked at four weeks likely due to technical factors. Specifically, a previous study reported that in large (>1 cm) glioma specimens, immunoreactivity of pERK was limited to the tumor periphery and was very low in the tumor core, as we also observed. This study suggested that due to time-dependent penetration of the formalin fixative, pERK state is lost in the deeper tissue cores by the time the fixative has permeated (Mandell, 2003).

The study of TNBC was extended then to demonstrate susceptibility of another TBNC cell line, MDA-MB-231. This cell line, which harbors an oncogenic mutations *KRAS^G13D^* and *BRAF^G464V^*, is characterized as *KRAS*-dependent (Scholl et al., 2009) and is highly sensitive to RRSP (Antic, 2015, Figure 4A, and Figure 5). In xenograft studies using MDA-MB-231, RRSP-DT_B_ induced a significant reduction in tumor size. Interestingly, the residual tumors did not show significant reduction in RAS or pERK, but instead revealed a patchy or focal distribution. Thus, the true reduction in RAS detection may be masked by the low vascularization of MDA-MB-231 tumors such that RRSP-DT_B_ reduces overall tumor size by diffusion from the tumor periphery rather than infiltrating equally throughout the tumor center, as occurred with the MDA-MB-436 xenografts.

We finally investigated the effects of RRSP-DT_B_ treatment on the CRC cell line HCT-116 (*KRAS^G13D^*). As with the other cell lines tested, RRSP-DT_B_ treatment resulted in complete RAS cleavage and pERK reduction in the low picomolar range, with a corresponding reduction in cell viability. Similar results were obtained in HCT-116 spheroids, including complete loss of RAS and pERK immunoreactivity. Notably, depletion of RAS and reduction in pERK were also observed in spheroids treated with the catalytically-inactive mutant RRSP*-DT_B_, but at a 10-fold higher dose. Previous binding studies showed that RRSP* retains its ability to bind recombinant KRAS, especially at high KRAS:RRSP* molar ratios (Biancucci et al., 2018). Therefore, although RRSP*-DT_B_ does not cleave RAS, it might impact its activity when used at high doses and after longer exposure both *in vitro* and *in vivo*.

As with the other tumor models, RRSP-DT_B_ treatment resulted in significant reductions in tumor size in an HCT-116 xenograft study. Both q.d. and b.i.d. dosing strategies showed similar median reductions in tumor size, however the b.i.d. arm showed less variability represented by a small interquartile range. Effective target engagement was demonstrated by a reduction in total RAS levels in residual RRSP-DT_B_ treated tumors. We did not observe statistically significant differences between controls and RRSP-DT_B_-treated tumors when analyzing pERK immunoreactivity. However, as noted above, large control tumors also showed poor immunostaining for pERK, and thus low values for control may have caused the lack of statistical significance in the small sample size when data are normalized. Even still, we observed reduced tumor size in all treatment groups and a decrease in pERK in the b.i.d. treatment group.

In order to assess the broader applicability of RRSP-DT_B_ and have a better understanding of its efficacy as an anti-cancer agent, RRSP-DT_B_ was tested against the NCI-60 cell line panel. Results from this large-scale screening showed that RRSP-DT_B_ inhibited cell growth in a wide variety of cell lines with diverse *RAS* genomic backgrounds and from different tissue types. Cell lines with genomic defects in *RAS* genes (e.g. missense mutations or amplifications) were enriched in the group highly responsive to RRSP-DT_B_. Moreover, we observed that although the expression levels of the DT receptor HB-EGF plays an important role in determining RRSP-DT_B_ susceptibility, some cancer cell lines that express less HB-EGF were still sensitive to RRSP-DT_B_, suggesting differential susceptibility of tumor cells to RAS/RAP1 inhibition. Thus, these data show that RRSP-DT_B_ has the potential to be broadly applicable against many types of cancer, including both wild-type and mutant *RAS* tumors. I addition, cancer cell lines from colon and lung are particularly sensitive to RRSP-DT_B_. However, several tumor cell lines with no genetic defects in RAS or the MAPK pathway remain very sensitive to RRSP-DT_B._

In total, this study provides solid proof-of-concept that the effective, receptor-mediated, intracellular delivery of a potent anti-RAS biologic represents a promising new approach for the development of RAS-targeted therapeutics. We contend that the ability of RRSP to directly and irreversibly inactivate both wild-type and mutant RAS proteins represents an attractive mechanism compared to the current approach of targeting RAS mutants individually. However, a pan-RAS inhibitor is expected to induce dose-limiting toxicity due to the critical importance of RAS signaling in non-cancerous tissues. The ability of the DT to deliver cargo exclusively to receptor-bearing cells provides a solution to this toxicity. While HB-EGF is upregulated on various tumor types, it is also widely expressed in humans, and may not represent the ideal receptor for tumor-targeting. In order to restrict the delivery of RRSP to tumor cells, the DT-based delivery platform described here can be re-targeted to various cell types by replacing the receptor-binding domain (DTR) of DT with other binding moieties, such as antibody fragments or ligands. The immunotoxin Ontak comprises wild-type diphtheria toxin with DTR replaced by interleukin 2 (IL-2), targeting Ontak to cells expressing the high-affinity IL-2 receptor (IL-2R) (Manoukian and Hagemeister, 2009). Similarly, RRSP-DT_B_ can be re-targeted in the same fashion, and RRSP-DTT-IL2 is able to efficiently cleave RAS in both MOLT-4 and Jurkat cell lines which express the IL-2R, but not in CFPAC-I cells, which does not (Figure S11). In fact, MOLT-4 was amongst the least responsive cell lines in the NCI-60 screen, suggesting that even cancers currently in the low sensitivity group would be much more sensitive to RRSP-DT_B_ following advanced engineering to target the chimeric toxin to alternate receptors appropriated for that cell type. Further study into engineered chimeric toxins, such as RRSP-DT_B,_ could usher in the next-generation of anti-cancer biologics.

## Materials and Methods

### Plasmids design, protein purification and endotoxin removal

DNA sequence corresponding to RRSP aa 3580-4089 was amplified from a plasmid containing the effector domain region cloned from *Vibrio vulnificus* CMCP6 in vector pXL PCR-TOPO (Kwak et al., 2011). The amplified gene was fused to different DT variants using the NEBuilder® HiFi DNA Assembly Cloning Kit (New England Biolabs Inc.). DNA sequence corresponding to CPD (nucleotides nt 12269-12894 of rtxA1) of MARTX toxin from *Vibrio vulnificus* was codon-optimized and synthesized as a double-stranded DNA fragment (IDT) and inserted into the DT vector (Auger et al., 2015) as above. A point mutation was made in RRSP using QuikChange Lightning Multi Site-Directed Mutagenesis Kit (Agilent Technologies) to change His4030 CAT codon to Ala (GCT) to generate the catalytically inactive RRSP_H4030A_ mutant. The final products were cloned into the Champion pET-SUMO expression system (Invitrogen). The different DT variants fused to RRSP were expressed as N-terminal His6-tagged and C-terminal StrepTagII-tagged proteins, transformed into either *E. coli* NiCo21(DE3) or BL21 (DE3) cells, and grown overnight in Luria-Bertani (LB) broth containing 50 µg/mL kanamycin. Next, overnight cultures were diluted 1:50 in fresh TB containing 35 µg/mL kanamycin and grown to OD_600_ = 0.8-1.0 at 37°C. Cultures were then induced with 1 mM isopropyl β-d−1-thiogalactopyranoside (IPTG) for 5 h at 25°C. Bacteria were pelleted by centrifugation at 10,000 rpm for 15 min and resuspended in 150 mL of Lysis Buffer (20 mM Tris-HCl pH 8.0, 500 mM NaCl, 20 mM imidazole, 2 mg/ml lysozyme and one tablet of EDTA-free protease inhibitor cocktail). Resuspended cells were sonicated using a Branson Digital Sonifier for 30 min (parameters: pulse ON 10 sec, pulse OFF 20 sec, 10 min, Amplitude 50%), the lysate spun down at 12,000 x g for 30 min and then loaded onto a 5 ml His-Trap Crude FF column (GE Healthcare) using a ÄKTA Purifier protein purification system (GE Healthcare). His-tagged proteins were eluted with a buffer containing 20 mM Tris-HCl pH 8.0, 500 mM NaCl and 500 mM imidazole. Fractions corresponding to the protein peaks were collected, pooled and loaded onto a gravity column containing Strep-Tactin Superflow high capacity resin (#2-1208-025, Iba Lifesciences). Strep-tagged proteins were eluted using 20 mM Tris-HCl pH 8.0, 150 mM NaCl and 10 mM d-Desthiobiotin. The His-Sumo tag was then removed by adding 0.01 µg of Sumo protease per 100 µg of purified protein in 20 mM Tris-HCl pH 8.0, 150 mM NaCl and 1 mM dithiothreitol at 30°C for 1 h. Next, proteins were further purified by size exclusion chromatography (SEC) using the HiLoad Superdex 16/600 200 prep grade column (GE Healthcare) run in 20 mM Tris-HCl pH 8.0 and 150 mM NaCl. Proteins were then dialyzed overnight using a ThermoFisher Slide-A-lyzer cutoff 20K in 20 mM Tris-HCl pH 8.0 and 150 mM NaCl and concentrated with a Millipore Amicon Ultra 30K spin concentrator. Finally, glycerol was added to the final protein solution containing 20 mM Tris-HCl pH 8.0, 150 mM NaCl, 10 mM imidazole and 8% glycerol. Protein purity was assessed by SDS-PAGE/Coomassie blue staining and concentration determined using NanoDrop ND1000 Spectrophotometer. Protein aliquots were flash frozen and stored at −80°C until use. For *in vivo* applications, endotoxin was removed from all protein preparations using Pierce High-Capacity Endotoxin Removal Resin (#88270) and residual endotoxin was quantified using Pierce LAL Chromogenic Endotoxin Quantitation Kit (#88282) following manufacturer’s instructions (Thermo Scientific).

### Cell lines and chemicals

All cell lines were purchased from the American Type Culture Collection, except for the *KRAS^WT^-*expressing RAS-less MEF cells that were kindly provided by the RAS Initiative at Frederick National Laboratory for Cancer Research (FNLCR) (Designation RPZ26216, expressed transgene KRAS 4B WT) (Drosten et al., 2010). Cells were cultured at 37°C and 5% CO_2_ atmosphere. MDA-MB-436, MDA-MB-231 and HCT-116 were grown in DMEM-F12 with Glutamax (Gibco) containing 10% fetal bovine serum (FBS; Gemini) and 1% penicillin/streptomycin (P/S; Invitrogen). Hs578T were grown in DMEM containing 10% FBS and 1% P/S (complete DMEM). MEF cells were cultured in complete DMEM with 4 µg/ml of blasticidin. All chemicals, unless otherwise specified, were purchased for Sigma-Aldrich.

### Antibodies

The anti-RAS 4E8 hybridoma cell line was kindly provided by the Frederick National Laboratory for Cancer Research (FNLCR). The antibody was purified by affinity chromatography as described in (Thomer et al., 2016). Antibody validation was performed as follows. 10 µM of recombinant K-, N- or H-RAS proteins were incubated alone or together with 1 µM of recombinant RRSP for 5 minutes at room temperature and then 1 µg of each sample was run on an SDS-PAGE gel followed by western blotting. The purified antibody used at 1:2000 dilution specifically recognized both cleaved and uncleaved bands of all three RAS isoforms (Figure S2I). This antibody thus is a pan-RAS monoclonal, here designated as mAb 4E8.

Some western blots and immunohistochemistry were performed with the commercially available anti-panRAS (Ras10, MA1-012; Thermo Fisher Scientific) antibody, that recognizes RAS Switch I and thus detects only uncleaved RAS. Other primary antibodies used are anti–Phospho-p44/42 MAPK (phosphorylated ERK1/2, Thr202/Tyr204, 197G2, Cell Signaling Technology #4377), anti-p44/42 MAPK (ERK1/2, L34F12, Cell Signaling Technology #4696), anti-HB-EGF (R&D Systems, #AF-259-NA;), and precision protein StrepTactin-HRP Conjugate rabbit Bio-Rad (#1610381) The anti-vinculin (Cell Signaling Technology #13901) was used for normalization. Secondary antibodies use were fluorescent-labeled IRDye 680RD goat anti-mouse (926-68070), IRDye 800CW goat anti-rabbit (925-322211) and IRDye 800CW donkey anti-goat (925-32214) from LI-COR Biosciences. Blot images were acquired using an Odyssey Infrared Imaging System (LI-COR Biosciences) and quantified by densitometry using NIH ImageJ software. Percentage of uncleaved RAS was calculated as previously described (Biancucci et al., 2018).

### SDS-PAGE and Western Blotting

At the experimental endpoint, cells were washed once with ice-cold PBS and protein lysates were extracted using M-PER mammalian protein extraction reagent (Thermo Fisher Scientific) with protease and phosphatase inhibitors (Sigma-Aldrich). Protein concentration was quantified using Bio-Rad protein assay dye reagent concentrate (#5000006). Equal amounts of proteins were separated by SDS-PAGE followed by western blot analysis as previously described (Biancucci et al., 2018).

### Cell viability, proliferation and growth inhibition assays

For quantitative viability assays, 10,000 cells per well were cultured in 96-well clear bottom white plates in the corresponding complete growth medium and treated the next day with RRSP-DT_B_ and RRSP*-DT_B_. At the end of treatments, CellTiter-Glo (Promega) was added to each well according to the manufacturer’s manual and luminescence was acquired using a Tecan Safire2 plate reader. CellTiter-Glo was incompatible with assessment of MDA-MB-436 cell viability under the experimental conditions used. Indeed, despite the cells being very sensitive to RRSP-DT_B_, no change in luminescence were detected between untreated and treated cells presumably due to release of ATP to the media that complicates the CellTiter-Glo reaction. Therefore, for this cell line, viability was determined via crystal violet assay. Briefly, MDA-MB-436 cells (10,000 cells/well) were plated in 96-well plates and treated with RRSP-DT_B_ and RRSP*-DT_B_. At the experimental endpoint, medium was carefully removed and cells gently washed with PBS. Crystal violet fixing/staining solution (0.05% (g/vol) crystal violet, 1% formaldehyde, 1% (v/v) methanol in phosphate buffered saline (PBS)) was then added to each well and the plate incubated at room temperature for 20 min. Next, wells were washed with tap water, air-dried, crystal violet solubilized using 10% acetic acid and absorbance recorded at 570 nm using a Tecan Safire2 plate reader. IC_50_ were calculated using the log(inhibitor) vs. response - variable slope (four parameters) function in Graphpad Prism. Crystal violet assay was also performed to visually assess viability of adherent cells using the same procedure described above but instead of solubilizing the dye, images of 6-well air-dried plates were acquired using a conventional desktop scanner. Cell proliferation was measured by colony-formation assay. Cells were treated with RRSP-DT_B_ and 10 nM RRSP*-DTB. After 72 hours, cells were harvested by trypsinization, counted on a hemocytometer and replated in 6-well plates at 2,500 cells per well. Colony formation was monitored over 10 days period, during which medium was replaced every two days. On day 10, crystal violet was applied to visualize the colonies. The open source ColonyArea ImageJ plug-in was used for quantitative analysis of the area % covered by the stained colonies (Guzman et al., 2014).

An NCI-60 five dose growth inhibition screen was performed on a panel of 60 human tumor cell lines derived from nine different tumor types by the National Cancer Institute Developmental Therapeutics Program (NCI-DTP) and growth inhibition percentage was calculated in accordance with their standard protocol previously published (Shoemaker, 2006). We normalized all data so that 0% growth inhibition corresponded to no change in the cell number following treatment and −100% or more was equal to cytotoxicity. Additional information about the screening methodology can be found on the NCI-DTP web site (https://dtp.cancer.gov/discovery_development/nci-60/methodology.htm).

### Maximum tolerated dose (MTD) and xenograft studies

The MTD study was performed at Charles River Laboratories (Wilmington, MA). Both female and male athymic NU(NCr)-*Foxn1^nu^* mice were used (5 mice/group). Mice received increasing dosing of RRSP-DT_B_ (0.004, 0.02, 0.1 and 0.5 mg/kg) and RRSP*-DT_B_ (0.5 mg/kg) via IP injection on an everyday schedule – weekends excluded – for two weeks. Mouse weight was monitored on a regular basis until the end of the experiment. Mice were humanely euthanized after loss of 20% body weight and counted as non-survivors.

MDA-MB-436 and MDA-MB-231 mouse xenografts were performed following our animal protocol, which was approved by the Institutional Animal Care and Use Committee (IACUC) at Northwestern University. Mice were maintained in Allentown cages in a sterile housing facility under controlled environmental conditions. 5-week old athymic NU(NCr)-*Foxn1^nu^* female mice were injected subcutaneously with 2.5×10^6^ cells in 100 µl/mouse of 50% 1X endotoxin-free PBS/50% Matrigel (Corning #356237) onto the right flank under anesthesia. On day 7, when tumors reached 100-200 mm^3^ in size, mice were randomized into groups of 5 and treatment started. For the MDA-MB-436 xenograft, control mice were administered IP with saline (endotoxin-free PBS) and treatment groups with 0.1 mg/kg of RRSP-DT_B_ and 0.1 mg/kg of RRSP*-DT_B_ on an every-other-day schedule (weekends excluded) for about 4 weeks. In a second experiment with the same cell line, mice were treated with 0.1 mg/kg of RRSP-DT_B_ every other day (EOD) and 0.1 mg/kg of RRSP-DT_B_ every day (ED) for about 4 weeks as well as with 0.25 mg/kg of RRSP-DT_B_ for two weeks followed by 0.1 mg/kg RRSP-DT_B_ for additional two weeks (weekends excluded). For the MDA-MB-231 xenograft, mice were treated with 0.1 mg/kg of RRSP-DT_B_ and 0.1 mg/kg of RRSP*-DT_B_ on an ED schedule (weekends excluded) for 4 weeks. Tumor size was measured twice a week using a digital caliper and tumor volume was calculated using the following formula: volume (mm^3^) = (l x w^2^)/2, where l is the length and w the width of the tumor. Mouse body weight was also measured twice a week.

The HCT-116 xenograft study was performed at Charles River Laboratories (Wilmington, MA). 5 x 10^6^ cells in 100 µl PBS (no Matrigel) were inoculated into the right flank of athymic NU(NCr)-*Foxn1^nu^* female mice. When tumors reached an average size of 80 - 120 mm^3^, mice were randomized into 5 groups of 10 and treatment started. The first group received PBS (saline), the second 0.1 mg/kg of RRSP-DT_B_ every day (1X/day), the third 0.1 mg/kg of RRSP-DT_B_ twice per day (2X/day), the fourth 0.1 mg/kg of RRSP*-DT_B_ 1X/day and the fifth 0.1 mg/kg of RRSP*-DT_B_ 2X /day. Both tumor size and mouse body weight were measured biweekly. For all the *in vivo* experiments performed in this study, when tumors exceeded 1500 mm^3^, mice were sacrificed as per protocol. At the end of the treatment schedule, mice were euthanized, tumors excised, and fixed in 10% formalin overnight.

### Spheroid formation, viability and image analysis

For tumor spheroids generation, a single cell suspension of HCT-116 cells was seeded at a concentration of 10,000 cells/well into 96-well ultra-low attachment plates (Corning #4520) in complete medium. Two days after cell seeding, treatments were added and viability assessed after 1, 3, 5 and 7 days using the Promega CellTiter-Glo 3D Cell Viability Assay following the manufacturer’s protocol. Spheroids were imaged at every time point with an EVOS XL Core imaging system and their volume analyzed using a freely available ImageJ plug-in as described in (Ivanov et al., 2014). Spheroids fixation and agarose-embedding prior to immunohistochemical analysis was performed as previously described (Vidimar et al., 2018).

### Immunohistochemistry of spheroids and *in vivo* tumors

Spheroids as well as *in vivo* tumors processing, paraffin-embedding, sectioning and immunohistochemical stainings were performed by the Robert H. Lurie Comprehensive Cancer Center’s Pathology Core Facility. Anti-rabbit pan-Ras (#PA5-85947; Thermo Fisher Scientific) and anti-rabbit Phospho-p44/42 MAPK (Erk1/2) (Thr202/Tyr204, (D13.14.4E) XP Cell Signaling #4370) antibodies were used at 1:500 and 1:300 dilutions, respectively. Primary antibodies were detected using a standard anti-rabbit secondary antibody followed by 3,3′-diaminobenzidine (DAB) revelation (Dako). Quantification of immunohistochemical signal intensity was performed by color deconvolution using ImageJ (Fiji version) as previously described (Vidimar et al., 2018).

### Bioinformatic analysis

Bioinformatics analyses were performed using R version 3.5.2. NCI-60 mutation data were retrieved from Reinhold WC et al (Reinhold et al., 2012). For the A549/ATCC and MDA-MB-231/ATCC cell lines, mutations were obtained from the ATCC website (https://www.atcc.org/~/media/PDFs/Culture%20Guides/Cell_Lines_by_Gene_Mutation.ashx). NCI-60 gene copy number alteration data and RNA expression z-scores were downloaded from cBioPortal (https://www.cbioportal.org) for the study with id “cellline_nci60” using the TCGAretriever R package (https://cran.r-project.org/web/packages/TCGAretriever/index.html). Bar plots, tile plots, and violin plots were built using ggplot2.

### Statistics

Statistical analysis was performed using Graphpad Prism software v.6. Bar plots represent mean of at least three independent experiments ± the standard deviation (SD). Statistical significance was determined using one-way analysis of variance (one-way ANOVA) assuming normal distribution. Dunnett’s multiple comparison post-test was used to compare the mean of treatment groups to the mean of the control group and Tukey’s multiple comparison test was used to compare the mean of each group with the mean of every other group. Points in the fitted dose-response curve are mean ± standard error of the mean (SEM). Statistical analysis on fold change data was carried out after log transformation of the data to obtain a more normalized distribution. For *in vivo* xenografts, data are reported as mean ± SEM and one-way ANOVA was performed to test for differences among the groups. Values of *p*<0.05 were considered statistically significant.

## Acknowledgments

We thank the Robert H. Lurie Comprehensive Cancer Center Pathology Core Facility and B. Shmaltuyeva, B. Frederick and Dr D. Gursel for assistance with immunohistochemical staining, A. Dean and C. Nordloh for assistance with xenografts, S. Son for performing validation of the pan-RAS antibody used in this study, Dr Damiano Fantini for bioinformatics assistance and the FNLCR for providing the RAS-less *KRAS^WT^* MEF cells and the anti-RAS 4E8 hybridoma cells.

## Funding

This work was funded by the Lynn Sage Cancer Research Foundation, Northwestern University Clinical and Translational Sciences (which is supported by NIH/NCATS Award UL1TR001422), the Northwestern Medicine Catalyst Fund, and the Robert H. Lurie Comprehensive Cancer Research Center (to K.S.). D.G. is supported by R01CA152601, R01CA152799, R01CA168292, R01CA214025, the Avon Breast Cancer Foundation, the Zell Family Foundation, the Chicago Biomedical Consortium, and the Searle Funds at The Chicago Community Trust. Additional support from the SickKids Proof-of-Principal Funding and the Canadian Institutes of Health Research 366017 (to R.M). M.B. was supported by PanCan/FNLCR Fellowship. G.B. was supported by the SickKids Restracomp Fellowship.

## Author contributions

V.V. designed and conducted all studies for TBNC and *in vitro* monolayer, spheroids, and immunohistochemistry for CRC. V.V. wrote the manuscript with editing provided by G.B., R.M. and K.S. M.P. designed, cloned, purified and initially tested RRSP fusions. Portions of this work were published as her Ph.D. dissertation. G.B. purified protein for CRO studies and conducted tests of RRSP fusions. M.B. contributed to the concept and design of RRSP-DT_B_. M.K. purified protein and assisted with TBNC mouse studies. D.G. contributed to design of TBNC animal studies. R.M. envisioned the concept of DT for delivery of cargo to cells, supervised engineering, testing of fusions, and design of CRO experiments. K.S. envisioned the concept of RRSP as a treatment for cancer and supervised all studies.

## Competing interests

M.B. and K.S. are authors of pending patent on use RRSP as a therapeutic for cancer (WO2016019379A1). K.S. is author on published patent on use of CPD for autoprocessing of proteins (US20100137563A1). R.M. and G.L.B. are authors on a published patent on use of DT for protein delivery and pending patent on RRSP-DT_B_ as a RAS-directed therapeutic (US20180080033A1 and WO2019104433A1, respectively). K.S. has a significant financial interest in Situ Biosciences LLC, which conducts contract research unrelated to this work.

## Data and materials availability

Plasmids for expression of RRSP-DT are available with materials transfer agreement from R.M., Hospital for Sick Children.

