## Supplemental Figures for "Inhibition of tumor growth by a novel engineered chimeric toxin that cleaves activated mutant and wild-type RAS"

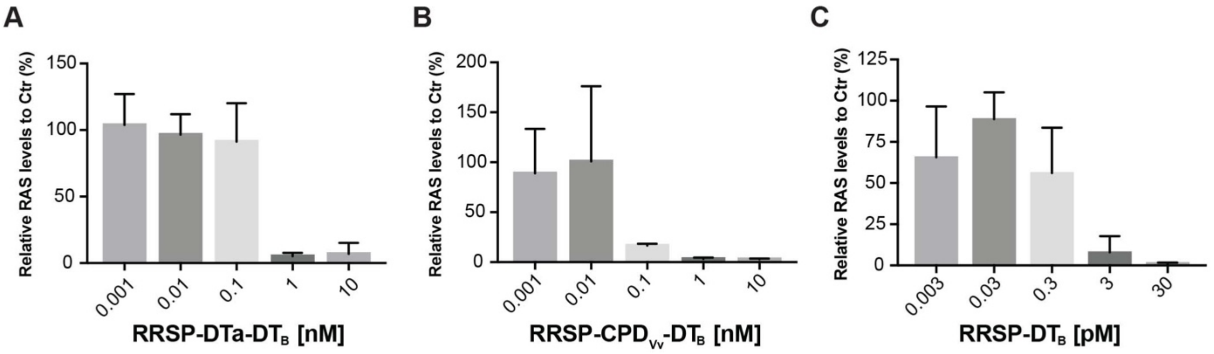


**Figure S1. Densitometry analysis showing the effect of various RRSP chimeras on total RAS levels in HCT-116 cells.** Densitometric analysis of western blots from HCT-116 cells (*KRAS*$^{G13D}$) treated with the indicated amount of various RRSP chimeras. (**A**) RRSP-DTa-DTB is RRSP fused to a detoxified mutant DT (K51E/E148K) via a (Gly-Gly-Gly-Gly-Ser)X2 ((G4S)2) linker. (**B**) RRSP-CPD$_{Vc}$-DTB is RRSP fused to the cysteine protease domain of the *V. vulnificus* MARTX toxin as in (A). The CPD is fused to DTB (diphtheria toxin residues 186-535) with the furin cleavage site mutated (RVRR->QVRQ) via a (G4S)2 linker. (**C**) RRSP-DTB is RRSP fused directly to DT residues 186-535 via a (G4S)2 linker.

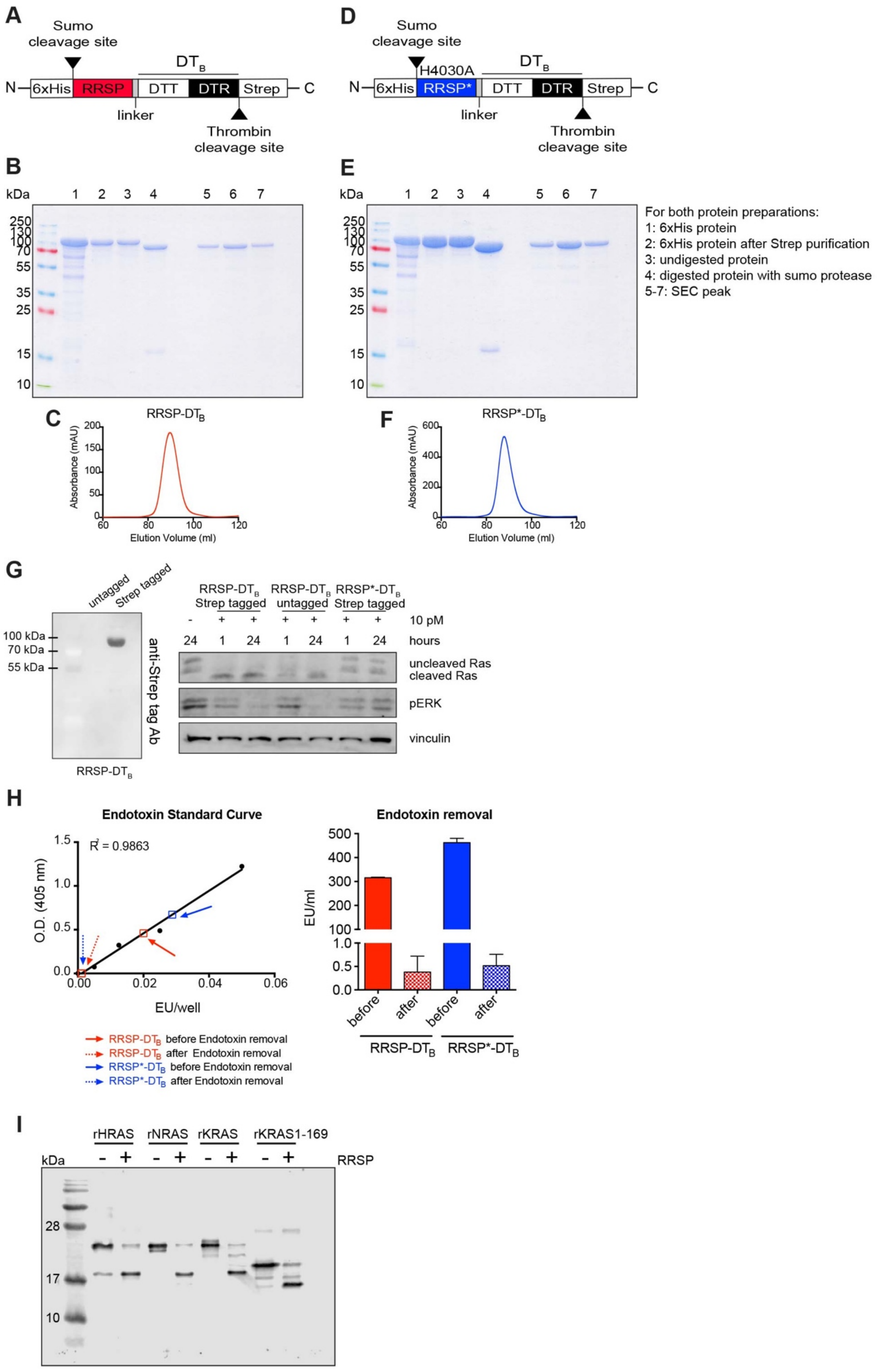
A
Sumo cleavage site
DT_B
N 6xHis RRSP DTT DTR Strep C
linker
Thrombin cleavage site
D
Sumo cleavage site
H4030A
N 6xHis RRSP* DTT DTR Strep C
linker
Thrombin cleavage site
B
E
kDa
250
130
100
70
55
35
25
15
10
For both protein preparations:
1: 6xHis protein
2: 6xHis protein after Strep purification
3: undigested protein
4: digested protein with sumo protease
5-7: SEC peak
C
RRSP-DT_B
Absorbance (mAU)
Elution Volume (ml)
F
RRSP*-DT_B
G
untagged
Strep tagged
100 kDa
70 kDa
55 kDa
anti-Strep tag Ab
RRSP-DT_B
RRSP-DT_B Strep tagged
RRSP-DT_B untagged
RRSP*-DT_B Strep tagged
10 pM
hours
uncleaved Ras
cleaved Ras
pERK
vinculin
H
Endotoxin Standard Curve
R² = 0.9863
O.D. (405 nm)
EU/well
RRSP-DT_B before Endotoxin removal
RRSP-DT_B after Endotoxin removal
RRSP*-DT_B before Endotoxin removal
RRSP*-DT_B after Endotoxin removal
Endotoxin removal
EU/ml
before
after
RRSP-DT_B
RRSP*-DT_B
I
rHRAS
rNRAS
rKRAS
rKRAS1-169
RRSP
kDa
28
17
10

Figure

**Figure S2. Purification of RRSP-DT$_B$ and RRSP*-DT$_B$ proteins and validation of panRAS antibody for detection of uncleaved/cleaved RAS. (A, D)** Schematic representation of RRSP-DT$_B$ (A) and RRSP*-DT$_B$ (D) plasmid constructs. **(B, E)**. Coomassie gel images of RRSP-DT$_B$ (B) and RRSP*-DT$_B$ (E) proteins showing degree of protein purity after nickel affinity purification, Strep-purification, removal of the SUMO-tag and size exclusion chromatography (SEC). (**C**, **F**). SEC peaks of purified RRSP-DT$_B$ (C) and RRSP*-DT$_B$ (F). **(G)** Western blot indicating presence of the Strep-tag in the RRSP-DT$_B$ purified protein. (**F**) Western blot showing the effect of Strep-tagged and untagged RRSP-DT$_B$ on RAS processing and phosphorylated ERK in HCT-116 cells. Cells were treated with 10 pM of RRSP-DT$_B$ for 1 and 24 hours. Strep-tagged RRSP*-DT$_B$ was also used as negative control. (**H**) Standard curve displaying efficiency in endotoxin removal from RRSP-DT$_B$ and RRSP*-DT$_B$ protein preparations for *in vivo* applications. Bar plots show residual endotoxin content (expressed in EU/ml) in both RRSP-DT$_B$ and RRSP*-DT$_B$ protein preparations. (**I**) Western blot image showing validation of the purified pan-RAS antibody used for detection of uncleaved/cleaved RAS. Recombinant HRAS, NRAS and KRAS were incubated with or without recombinant RRSP and samples were run on a SDS-PAGE gel followed by western blotting. The antibody recognized both cleaved and uncleaved bands of all RAS isoforms.

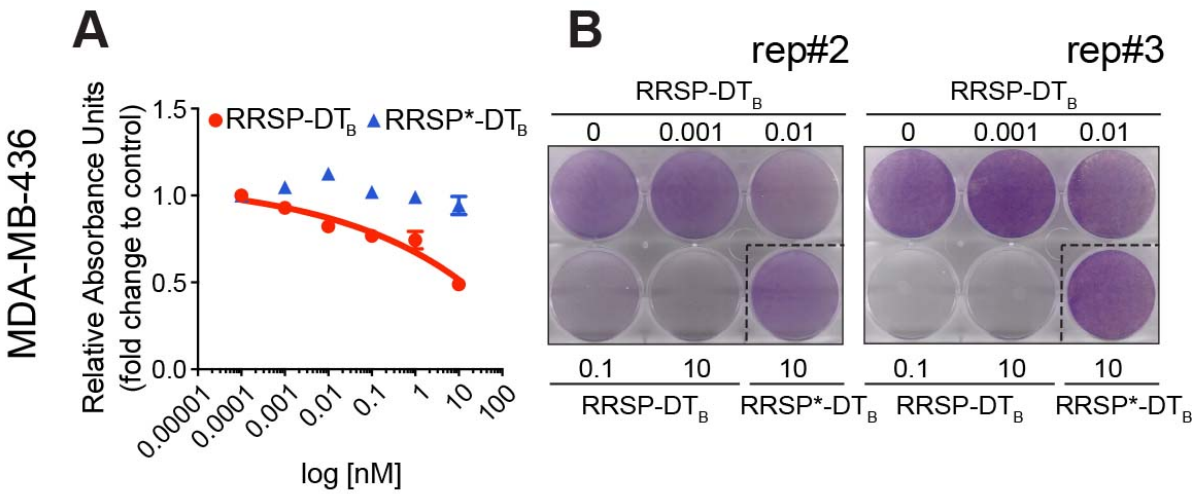


**Figure S3. Additional data on the effect of RRSP-DT$_B$ on viability of MDA-MB-436.** (**A**). Dose-response curve of RRSP-DT$_B$ in MDA-MB-436 cells following 24 h of treatment. Slopes of the dose-response curves for both RRSP-DT$_B$ and mutant RRSP*-DT$_B$ were not steep enough to retrieve an $IC_{50}$. Results are expressed as means ± SEM ($n$=3). (**B**) Additional two replicate images of crystal violet staining of MDA-MB-436 cells treated with RRSP-DT$_B$ and RRSP*-DT$_B$ as indicated for 72 h.

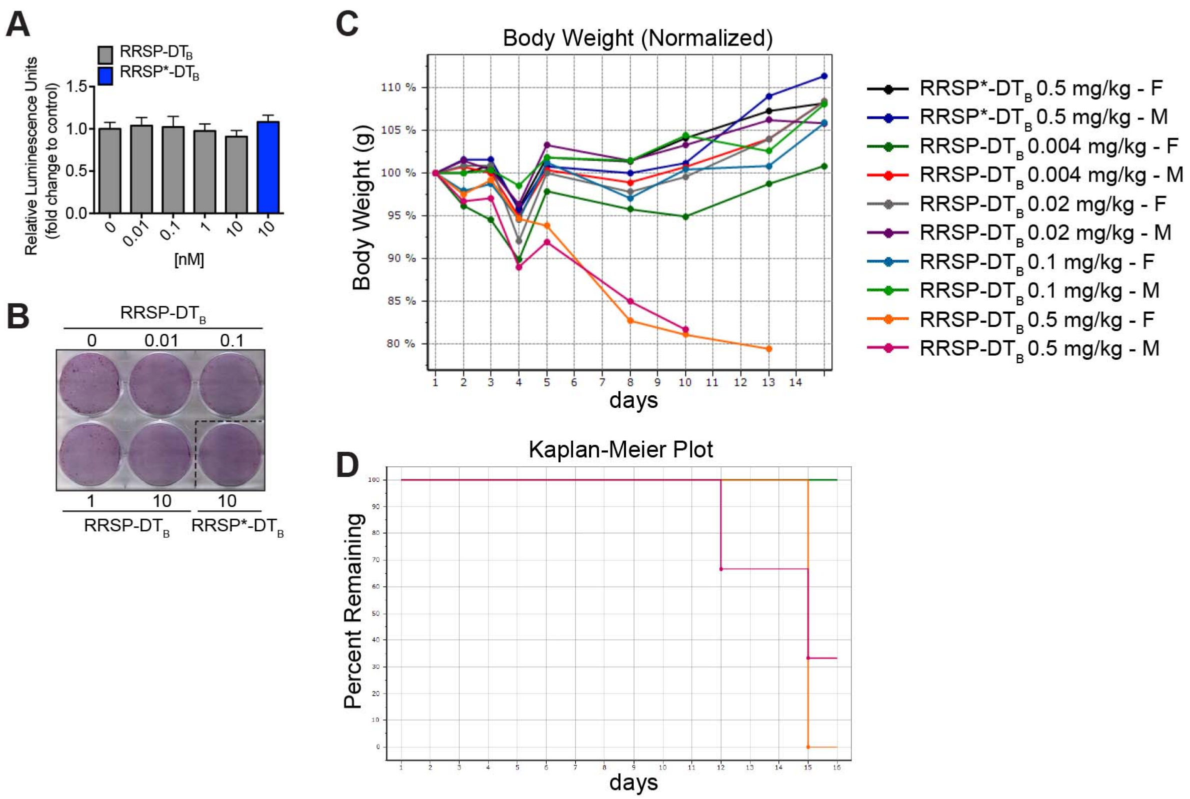


**Figure S4. Effect of RRSP-DTB on mouse embryonic fibroblasts (MEF) and *in vivo* Maximum Tolerated Dose (MTD) study.** (**A**) Viability and (**B**) crystal violet staining of RAS-less $KRAS^{WT}$ MEF cells following 72 h of treatment with RRSP-DTB and RRSP*-DTB as indicated. (**C**) Body weight expressed as percentage of initial weight over time from the MTD study that was performed treating athymic female *nu/nu* nude mice at the indicated doses every day for two weeks (weekends excluded). (**D**) Kaplan-Maier plot showing overall survival of mice treated with RRSP-DTB and RRSP*-DTB.

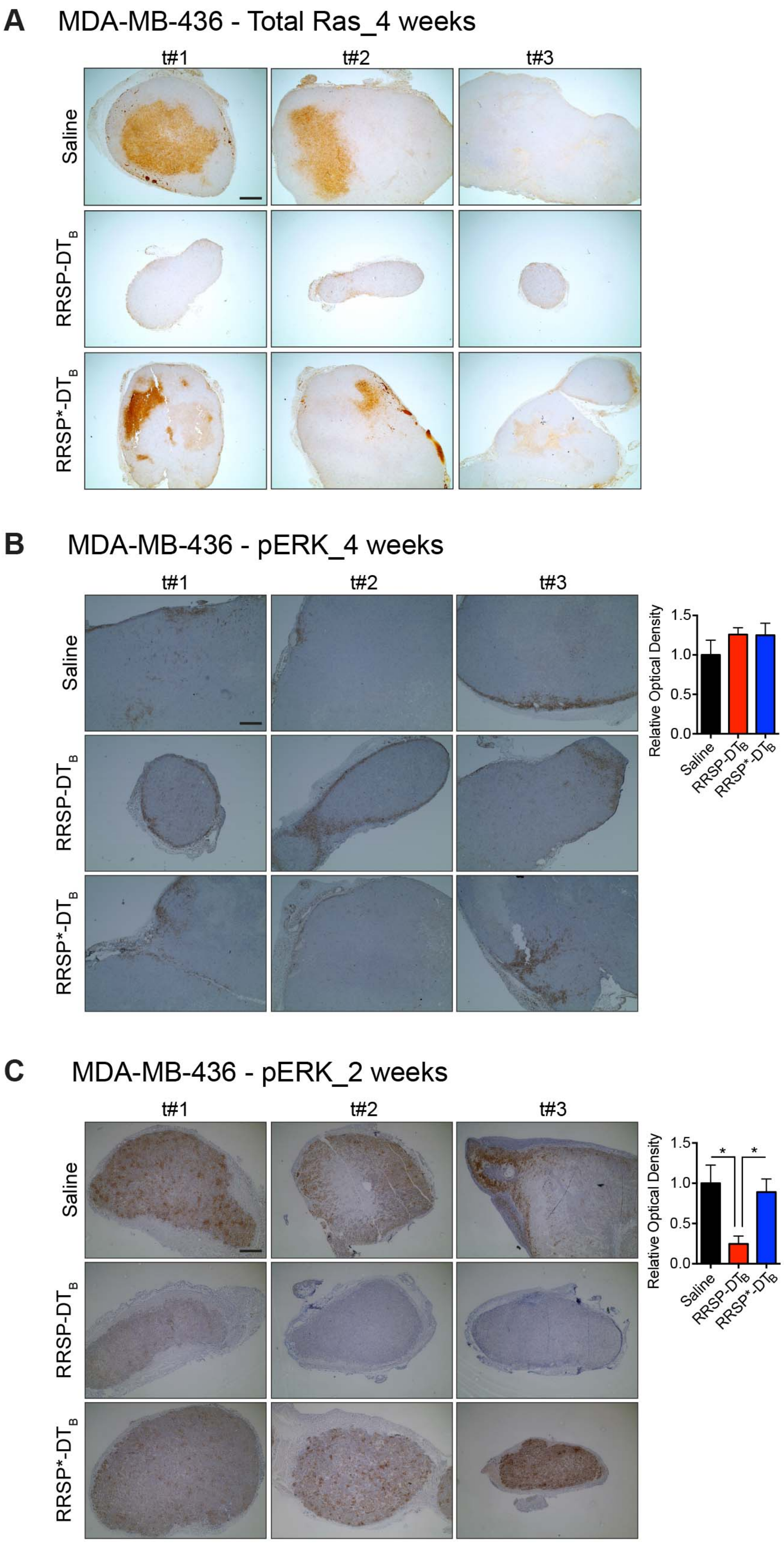
A
MDA-MB-436 - Total Ras_4 weeks
t#1
t#2
t#3
Saline
RRSP-DTB
RRSP*-DTB
B
MDA-MB-436 - pERK_4 weeks
t#1
t#2
t#3
Saline
RRSP-DTB
RRSP*-DTB
Relative Optical Density
1.5
1.0
0.5
0.0
Saline
RRSP-DTB
RRSP*-DTB
C
MDA-MB-436 - pERK_2 weeks
t#1
t#2
t#3
Saline
RRSP-DTB
RRSP*-DTB
Relative Optical Density
1.5
1.0
0.5
0.0
*
*
Saline
RRSP-DTB
RRSP*-DTB

**Figure S5. Whole tumor IHC images and analysis of total RAS and pERK levels from the MDA-MB-436 xenograft.** (**A**) Sections of whole tumors from the MDA-MB-436 xenograft immunostained with a total RAS antibody ($n$ = 3; scale bar = 200 μm). Mice were treated every other day (weekends excluded) at the indicated treatment conditions for about 4 weeks. (**B**) Sections of whole tumors from the same MDA-MB-436 xenograft immunostained with a pERK antibody (scale bar = 100 μm). Bar plot shows quantification of pERK DAB signal via color deconvolution ($n$ = 3). (**C**). Sections of whole tumors from a shorter, 2-week long MDA-MB-436 xenograft immunostained with a pERK antibody and quantification of DAB signal via color deconvolution (*$p < 0.05$, one-way ANOVA followed by Tukey's multiple comparison test, $n$=3; scale bar = 200 μM).

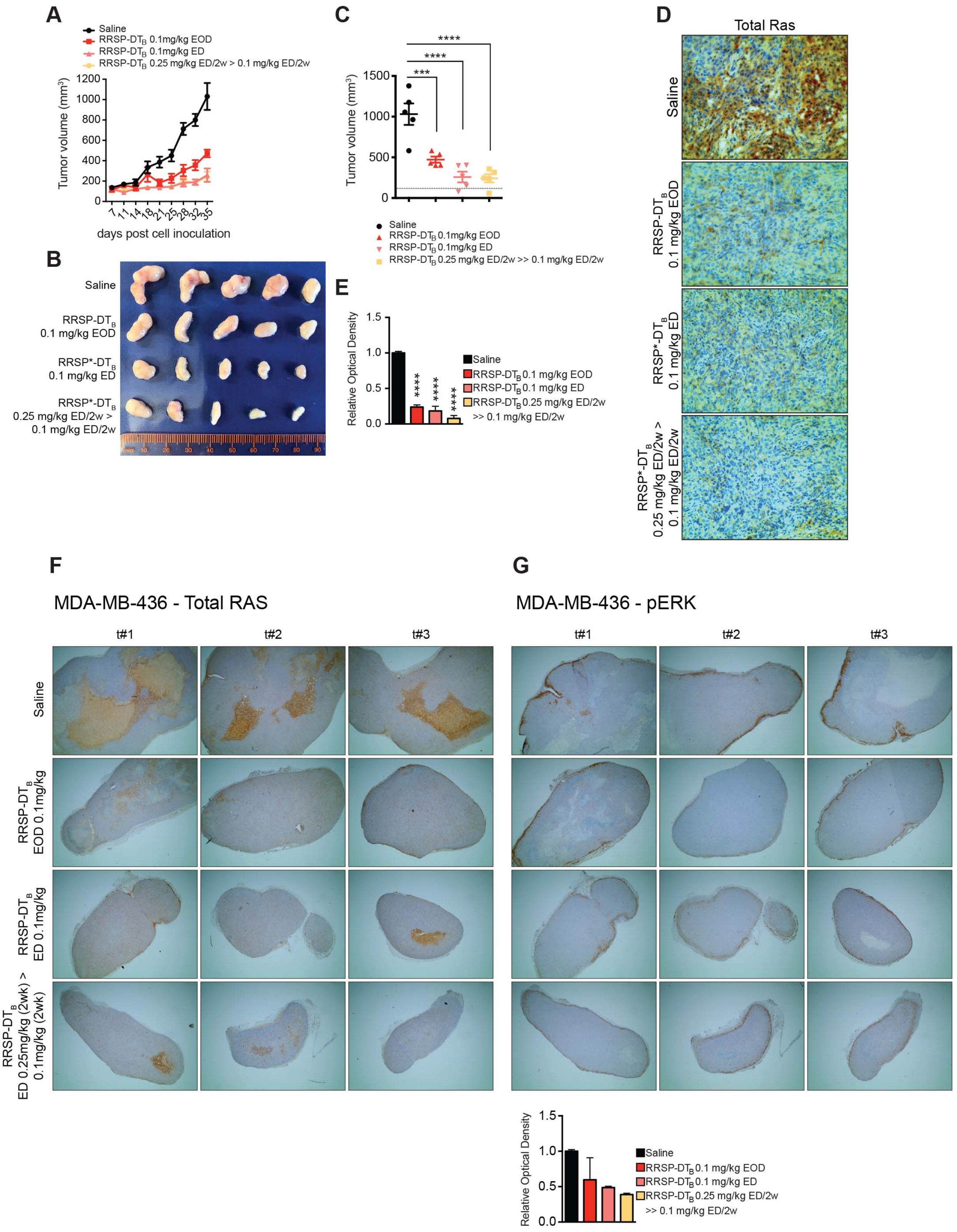
A
Saline
RRSP-DTB 0.1mg/kg EOD
RRSP-DTB 0.1mg/kg ED
RRSP-DTB 0.25 mg/kg ED/2w > 0.1 mg/kg ED/2w
Tumor volume (mm3)
days post cell inoculation
B
Saline
RRSP-DTB 0.1 mg/kg EOD
RRSP*-DTB 0.1 mg/kg ED
RRSP*-DTB 0.25 mg/kg ED/2w > 0.1 mg/kg ED/2w
C
Tumor volume (mm3)
Saline
RRSP-DTB 0.1mg/kg EOD
RRSP-DTB 0.1mg/kg ED
RRSP-DTB 0.25 mg/kg ED/2w >> 0.1 mg/kg ED/2w
D
Total Ras
Saline
RRSP-DTB 0.1 mg/kg EOD
RRSP*-DTB 0.1 mg/kg ED
RRSP*-DTB 0.25 mg/kg ED/2w > 0.1 mg/kg ED/2w
E
Relative Optical Density
Saline
RRSP-DTB 0.1 mg/kg EOD
RRSP-DTB 0.1 mg/kg ED
RRSP-DTB 0.25 mg/kg ED/2w >> 0.1 mg/kg ED/2w
F
MDA-MB-436 - Total RAS
t#1
t#2
t#3
Saline
RRSP-DTB EOD 0.1mg/kg
RRSP-DTB ED 0.1mg/kg
RRSP-DTB ED 0.25mg/kg (2wk) > 0.1mg/kg (2wk)
G
MDA-MB-436 - pERK
t#1
t#2
t#3
Relative Optical Density
Saline
RRSP-DTB 0.1 mg/kg EOD
RRSP-DTB 0.1 mg/kg ED
RRSP-DTB 0.25 mg/kg ED/2w >> 0.1 mg/kg ED/2w

**Figure S6. Summary of the effect of RRSP-DT$_B$ on an additional MDA-MB-436 xenograft.** (**A**) Tumor growth curve of vehicle and RRSP-DT$_B$-treated athymic *nu/nu* female mice bearing MDA-MB-436 tumors at the indicated doses and treatment schedule. EOD, every other day. ED, every day. (**B**) Representative images of MDA-MB-436 tumors at the experimental endpoint. (**C**) Waterfall plots indicating the change in percentage tumor volume compared to initial tumor volume. Each bar represents an individual tumor. (**D**) Column scatter plots showing individual tumor volumes at the end of the treatment schedule. Horizontal dashed lines indicate the baseline tumor volume on the first day of treatment, which corresponds to the average of tumor volumes at the indicated time (118 $mm^3$). Data are means ± SEM and *n*=5 mice in every group. Statistical analysis between vehicle and treatment groups was performed using one-way ANOVA followed by Tukey's multiple comparison test (****p* < 0.001, *****p* < 0.0001). (**E**). Representative IHC images of immunoreactivity to total RAS in sections from MDA-MB-436 tumors and corresponding quantification of DAB optical density via color deconvolution using whole tumor sections (*****p* < 0.0001, one-way ANOVA followed by Tukey's multiple comparison test, *n*=3; scale bar = 100 μM). (**F**). Sections of whole tumors from the MDA-MB-436 xenograft immunostained for a total RAS antibody and (**G**) ERK phosphorylation (*n* = 3; scale bar = 200 μm). Bar plot shows quantification of pERK DAB signal via color deconvolution.

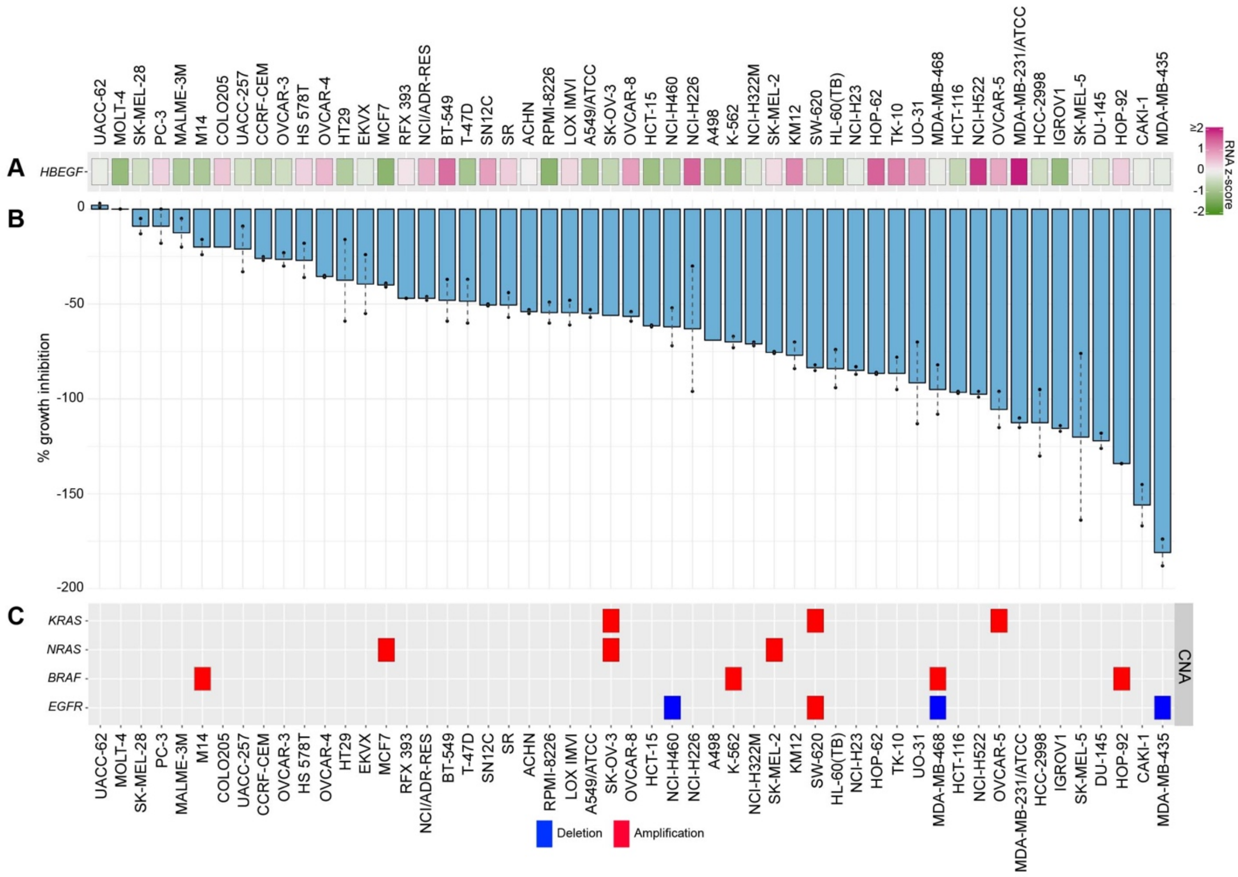


**Figure S7. Expression of HB-EGF and copy number alterations of *KRAS, NRAS, BRAF* and *EGFR* across several cell lines from the NCI-60 panel.** (**A**). Heatmap of RNA-Seq expression z-scores for human HB-EGF in 53 out of 60 cell lines included in the NCI-60 panel. Data were retrieved from cBioPortal (https://www.cbioportal.org) for the study with id "cellline_nci60". The color key indicates gene expression value (pink for upregulated and green for downregulated). (**B**). Bar plot showing the percent growth inhibition of RRSP-DT$_B$ for the 56 cell lines included in the analysis. Cancer cell lines were ranked in descending order based on their growth inhibition % value. The presence of two dots on the bars indicate that two replicates were performed per each cell line and bars represent means. No dots indicate that only one replicate was available. (**C**). Map of copy number alterations in *KRAS, NRAS, BRAF* and *EGFR* genes across the 53 cell lines from the NCI-60 panel. Red indicates gene amplification and blue gene deletions.

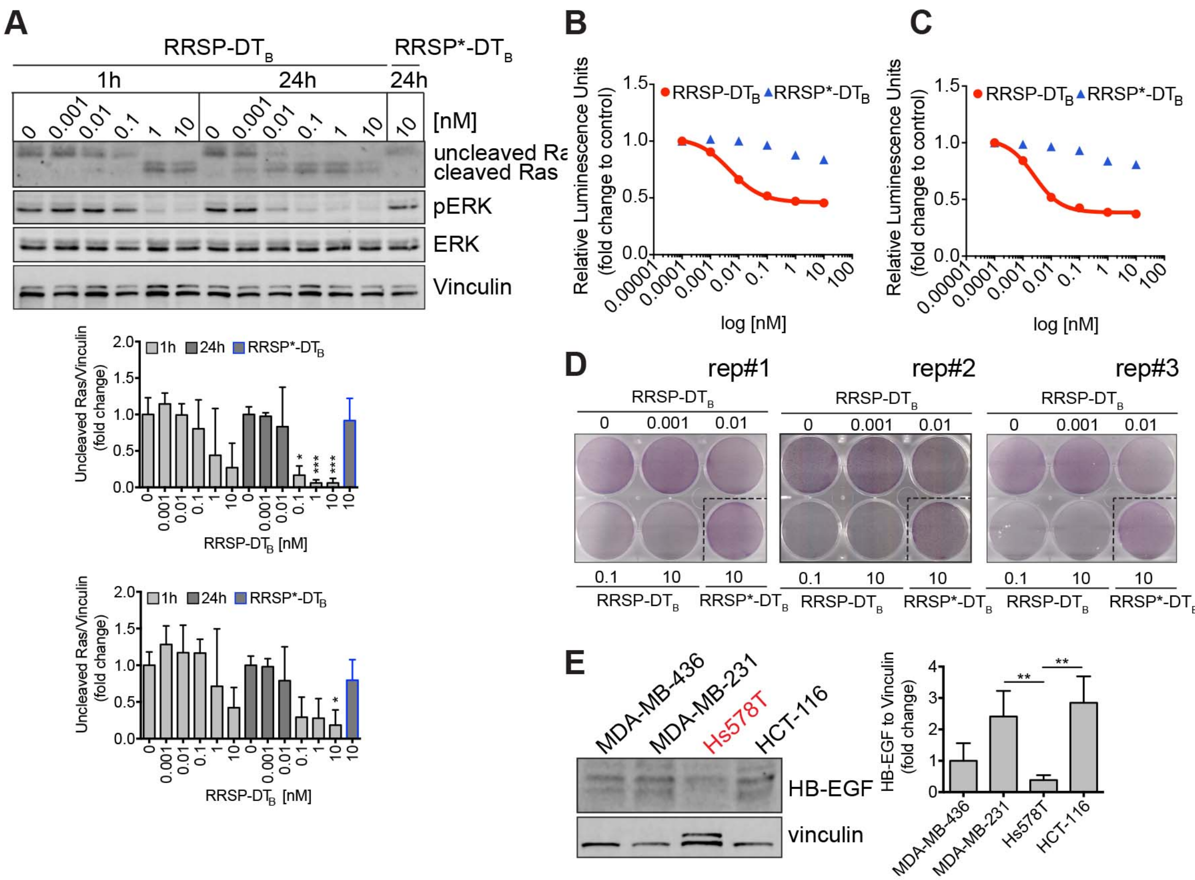


**Figure S8. Effect of RRSP-DT$_B$ on the TNBC Hs578T *HRAS G12D* cell line.** (**A**). Representative western blot and densitometric analysis of uncleaved RAS and phosphorylated ERK in Hs578T *HRAS G12D* cells treated with increasing doses of RRSP-DT$_B$ for 1 and 24 h. The catalytically-inactive RRSP*-DT$_B$ mutant was used as negative control at 10 nM and vinculin as gel loading control. Results are expressed as means ± SD of three independent experiments (*$p < 0.05$, ***$p < 0.001$ versus corresponding control 0 nM; one-way ANOVA followed by Dunnett's multiple comparison test, n = 3). (**B** and **C**) Fitted dose-response curve of RRSP-DT$_B$ in Hs578T cells following 24 (B) and (C) 72 h of treatment. Slopes of the dose-response curves for both RRSP-DT$_B$ and mutant RRSP*-DT$_B$ were not steep enough to retrieve an $IC_{50}$. Results are expressed as means ± SEM ($n$=3). (**D**). Images of crystal violet-stained plates in triplicate of Hs578T cells treated with RRSP-DT$_B$ and RRSP*-DT$_B$ as indicated for 72 h. (**E**). Western blot image showing expression of the DT receptor HB-EGF across MDA-MB-436, MDA-MB-231, Hs578T and HCT-116 cell lines.

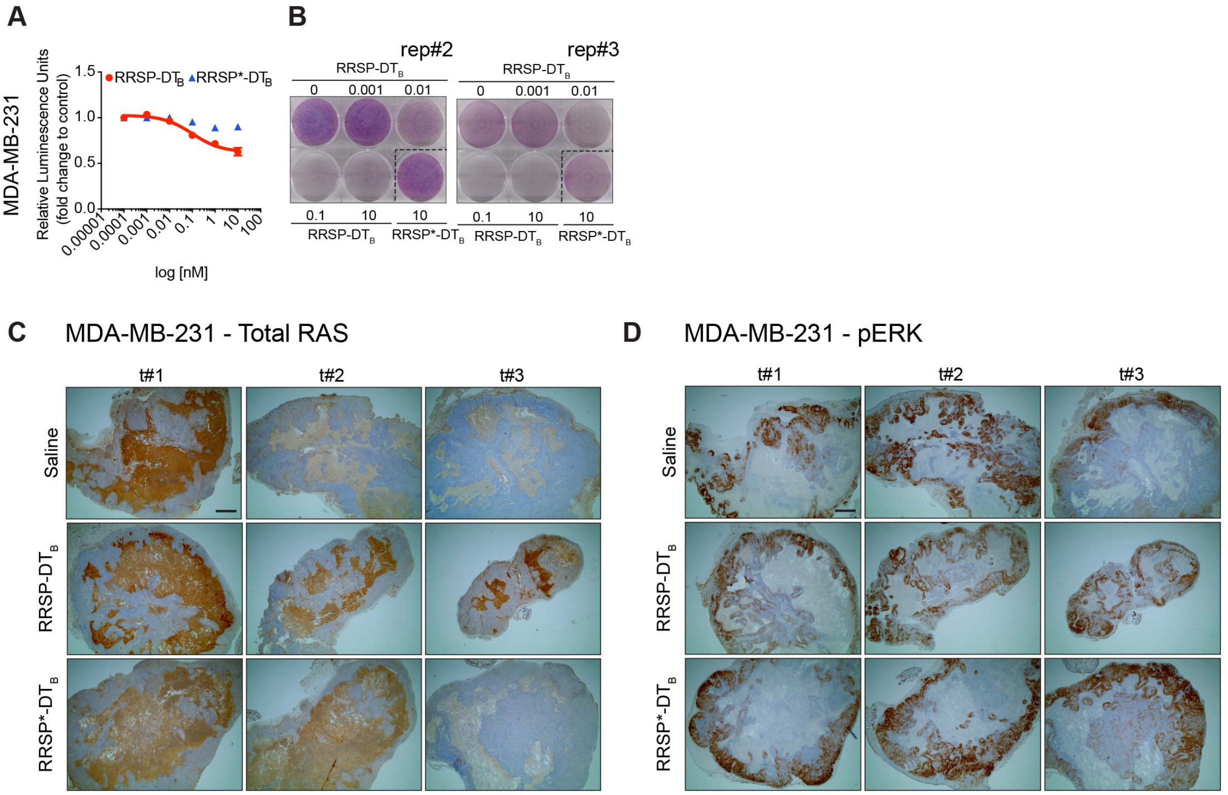


**Figure S9. Additional data on the effect of RRSP-DT$_B$ on viability of MDA-MB-231 cells and whole tumor IHC images and analysis of total RAS and phosphorylated ERK from the MDA-MB-231 xenograft.** (**A**). Dose-response curve of RRSP-DT$_B$ in MDA-MB-231 cells following 24 h of treatment. Slopes of the dose-response curves for both RRSP-DT$_B$ and mutant RRSP*-DT$_B$ were not steep enough to retrieve an $IC_{50}$. Results are expressed as means ± SEM (*n*=3). (**B**) Additional two replicate images of crystal violet staining of MDA-MB-231 cells treated with RRSP-DT$_B$ and RRSP*-DT$_B$ as indicated for 72 h. (**C**) Sections of whole tumors from the MDA-MB-231 xenograft immunostained with a total RAS antibody (n = 3; scale bar = 200 µm). Mice were treated every day (weekends excluded) at the indicated treatment conditions for about 4 weeks. (**D**) Sections of whole tumors from the same MDA-MB-231 xenograft immunostained with a pERK antibody (scale bar = 200 μm). Bar plot shows quantification of total RAS and phosphorylated ERK DAB signal using color deconvolution ($n$ = 3).

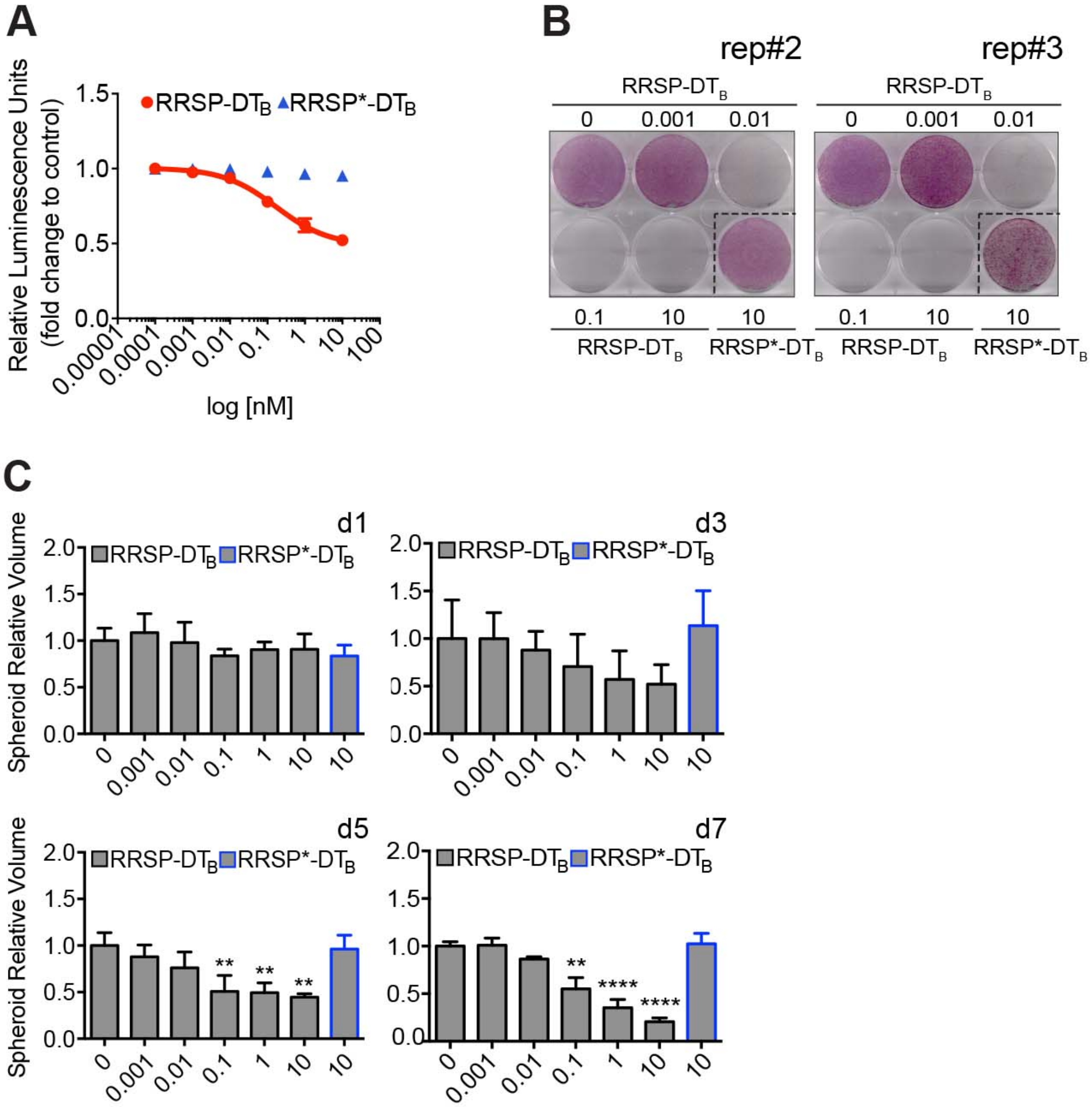


**Figure S10. Effect of RRSP-DT$_B$ on viability of CRC HCT-116 *KRAS G13D* cells after 24 hours of treatment and spheroids' volume quantification.** (**A**) Dose-response curve of RRSP-DT$_B$ in HCT-116 cells following 24 h of treatment. Slopes of the dose-response curves for both

RRSP-$DT_B$ and mutant RRSP*-$DT_B$ were not steep enough to retrieve an $IC_{50}$. Results are expressed as means ± SEM (*n*=3). (**B**) Additional two replicate images of crystal violet staining of HCT-116 cells treated with 0 RRSP-$DT_B$ or RRSP*-$DT_B$ as indicated for 72 h. (**C**). Quantitative analysis of spheroids' volumes. Results are expressed as means ± SD of three independent experiments (**$p < 0.01$, ****$p < 0.0001$ versus corresponding control 0 nM; one-way ANOVA followed by Dunnett's multiple comparison test, n = 3).

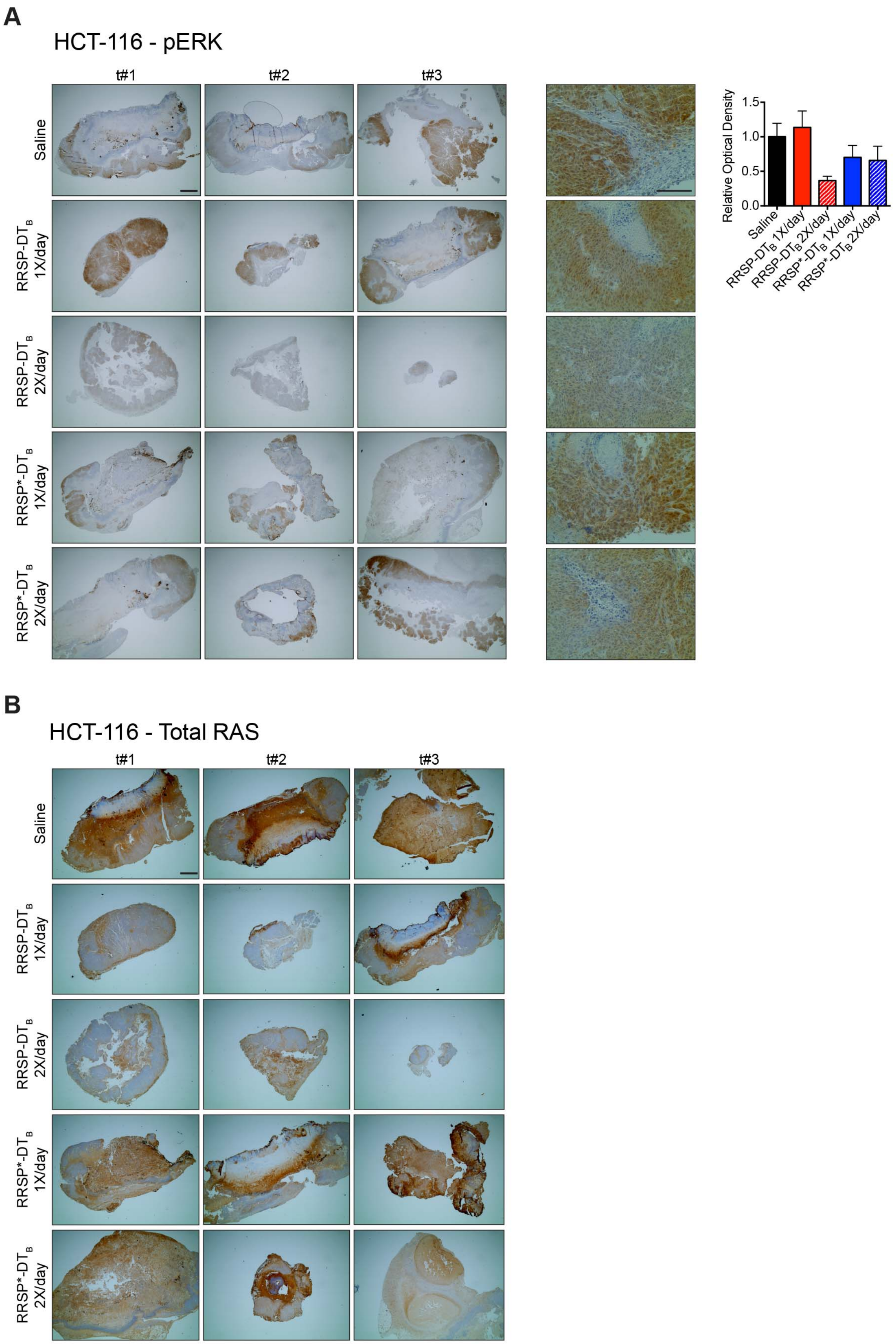
A
HCT-116 - pERK
t#1
t#2
t#3
Saline
RRSP-DT$_B$ 1X/day
RRSP-DT$_B$ 2X/day
RRSP*-DT$_B$ 1X/day
RRSP*-DT$_B$ 2X/day
Relative Optical Density
1.5
1.0
0.5
0.0
B
HCT-116 - Total RAS

**Figure S11. Whole tumor IHC images and analysis of total RAS and phosphorylated ERK from the HCT-116 xenograft.** (**A**) Sections of whole tumors from the HCT-116 xenograft immunostained with a total RAS antibody (*n* = 3; scale bar = 200 µm). Mice were treated every day (1X/day) and twice per day (2X/day) weekends excluded at the indicated treatment conditions for about 4 weeks. (**B**) Sections of whole tumors from the same HCT-116 xenograft immunostained for phosphorylated ERK (scale bar = 200 µm). Bar plot shows quantification of phosphorylated ERK DAB signal using color deconvolution (*n* = 3).

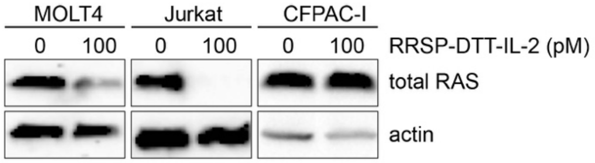


**Figure S12. Selective targeting of RRSP-DT$_B$ to IL-2 expressing cancer cells via replacement of the DTR domain.** Western blot image showing total RAS levels following treatment with RRSP-DTT-IL-2 of lymphoblast MOLT4 and Jurkat cells expressing high levels of IL-2 receptor and pancreatic CFPAC-I cells expressing none to low levels of the IL-2 receptor.
